# Follicular helper T cells expressing Blimp1 are specialized for plasma cell differentiation

**DOI:** 10.1101/2022.03.31.486642

**Authors:** Karen B. Miller, Andrew G. Shuparski, Brett W. Higgins, Siddhartha M. Sharma, Pierre J. Milpied, Louise J. McHeyzer-Williams, Michael G. McHeyzer-Williams

## Abstract

B cells differentiate into antibody-producing plasma cells (PC) and germinal center (GC) B cells under the guidance of specialized CD4^+^ follicular helper T (T_FH_) cells. Here, we demonstrate that CD4 T cells require *Prdm1* expression for both early PC differentiation and post-GC PC formation. Using dual Blimp1/Foxp3 reporter mice and single cell-indexed analysis, we segregate persistent compartments and expressed transcriptional programs of Blimp1^+^ CXCR5^+^PD1^hi^ T_FH_ (referred to here as PC-T_FH_) from canonical Blimp1^-^ *Bcl6*^+^ T_FH_ (GC-T_FH_) and Blimp1^+^Foxp3^+^ T_FR_ immune regulators. Antigen recall expands localized PC-T_FH_ compartments with rapidly divergent antigen-specific memory PC-T_FH_ and GC-T_FH_ programs. Thus, Blimp1 is a central mediator of PC-T_FH_ function producing specialized T_FH_ subsets that co-ordinate with GC-T_FH_ function to establish high-affinity long-lasting protective immunity to vaccines and infection.

**One-Sentence Summary:** Blimp1 expressing T_FH_ cells express unique transcriptional programs to control PC formation

**RESEARCH ARTICLE SUMMARY:** *Introduction:* Adaptive B cell immunity rapidly emerges to form plasma cells (PC) for antibody production and non-PC that enter germinal centers (GC) to evolve higher affinity B cell receptors. Both pathways are essential to long-term high-affinity immune protection. The early PC to GC cell fate division is driven by B cell expression of mutually antagonist transcriptional repressors Blimp1 and Bcl6. This dichotomous B cell outcome is orchestrated through antigen-specific contact by follicular helper T (T_FH_) cells that express Bcl6 to upregulate CXCR5, localize into B cell regions and express transcriptional programs that influence B cell fate and function. It remains unclear what T_FH_ cell mechanisms differentially impact these divergent B cell pathways.

*Rationale:* Blimp1 is found in Foxp3^+^ follicular regulatory T (T_FR_) cells known to impact GC B cell outcomes and play a role controlling antibody-mediated autoimmunity. In the context of infection, induced Blimp1 expression in CD4 T cells is expressed by conventional non-T_FH_ effector cell compartments. Blimp1 segregates with emigrant CD4 T cells that leave the reactive lymphoid tissue to control innate immune function at the site of antigen entry. Conversely, Bcl6 is predominantly expressed in the GC regulating T_FH_ pathway and is demonstrated to suppress Blimp1 expression. Germline ablation of Bcl6 exaggerates type 2 effector T_H_ cell functions that promote excessive antibody production in the absence of the GC reaction. Similarly, loss of Bcl6 in CD4 T cells abrogates GC formation and post-GC PC responses, however multiple recent reports indicate continued support for antibody production without a Bcl6^+^ T_FH_ compartment. To reconcile these findings, we propose a division of T_FH_ function with separable pathways to regulate PC and GC differentiation. We hypothesize a central role for persistent CD4 T cell expressed Blimp1 that segregates early T_FH_ transcriptional control to create an effector cell program that selectively targets PC differentiation.

*Results:* Direct intracellular staining for protein, confirmed with single Blimp1 and dual (Foxp3) reporter mice, identified Blimp1 expressing CXCR5^+^PD1^hi^ T_FH_ and T_FR_ subsets within the spleen, bone marrow and other lymphoid tissues at steady-state. Conditional deletion of *Prdm1* in CD4 T cells and adoptive transfer into immunodeficient hosts with splenic B cells, truncated both early pre-GC and late post-GC formation of PC providing a causal link to both pathways of differentiation in vivo. Across steady-state splenic T cells, in vitro activated Blimp1^+^CD25^-^ CD4 T cells in T-B cell co-cultures correlated with significant levels of PC induction. Integrated single cell-indexed strategies segregate the transcriptional programs of Blimp1 expressing T_FH_ cells (referred to here as PC-T_FH_) from canonical GC-inducing Bcl6^+^ T_FH_ cells (GC-T_FH_), both distinct from Blimp1^+^ T_FR_ cell programs in the steady-state. Immunization and recall produce follicular localized PC-T_FH_ with pMHCII-tetramer binding memory response T_FH_ cells that segregate across PC-T_FH_ and GC-T_FH_ compartments re-iterating the dichotomous transcriptome seen at steady-state.

*Conclusion:* This study identifies Blimp1 as a key mediator of PC-T_FH_ cells that sub-specialize as inducers of PC differentiation and bifurcate from the Bcl6^+^ GC-T_FH_ cell pathway and functions. Persistent PC-T_FH_ compartments assort across multiple lymphoid tissues at steady-state and are distinct from Foxp3^+^Blimp1^+^ T_FR_ immune regulators. While PC T_FH_ cells alone are required for early and rapid antibody responses, both T_FH_ sub-classes are essential to the generation of high-affinity long-lived and memory response PC compartments. Cellular organization and molecular components of the PC-T_FH_ transcriptional program indicate functional sub-specialization that can be separately targeted for immunotherapeutic purposes and adjuvant design in future vaccines.

*Sub-specialized Blimp1^+^ PC-T_FH_ cells control PC differentiation:* Adaptive immune protection requires balancing the evolution of BCR affinity within germinal center (GC) B cells and the differentiation of plasma cells (PC) for production of antibodies. Both functional B cell pathways require the antigen-specific induction of specialized CD4^+^ follicular T (T_FH_) cells. Within GC-inducing T_FH_ cells, Bcl6 is required to drive the formation and function of GC B cells. Here, we segregate PC-inducing T_FH_ cells that require Blimp1 as a key mediator of antigen-specific PC differentiation. The Blimp1^+^ PC-T_FH_ transcriptional program diverges from Bcl6^+^ GC-T_FH_ compartment and Blimp1^+^Foxp3^+^ follicular regulatory T (T_FR_) compartments. Antigen-specific PC-T_FH_ emerge and segregate rapidly from GC-T_FH_ after priming and recall to co-operatively induce effective long-term adaptive immunity. 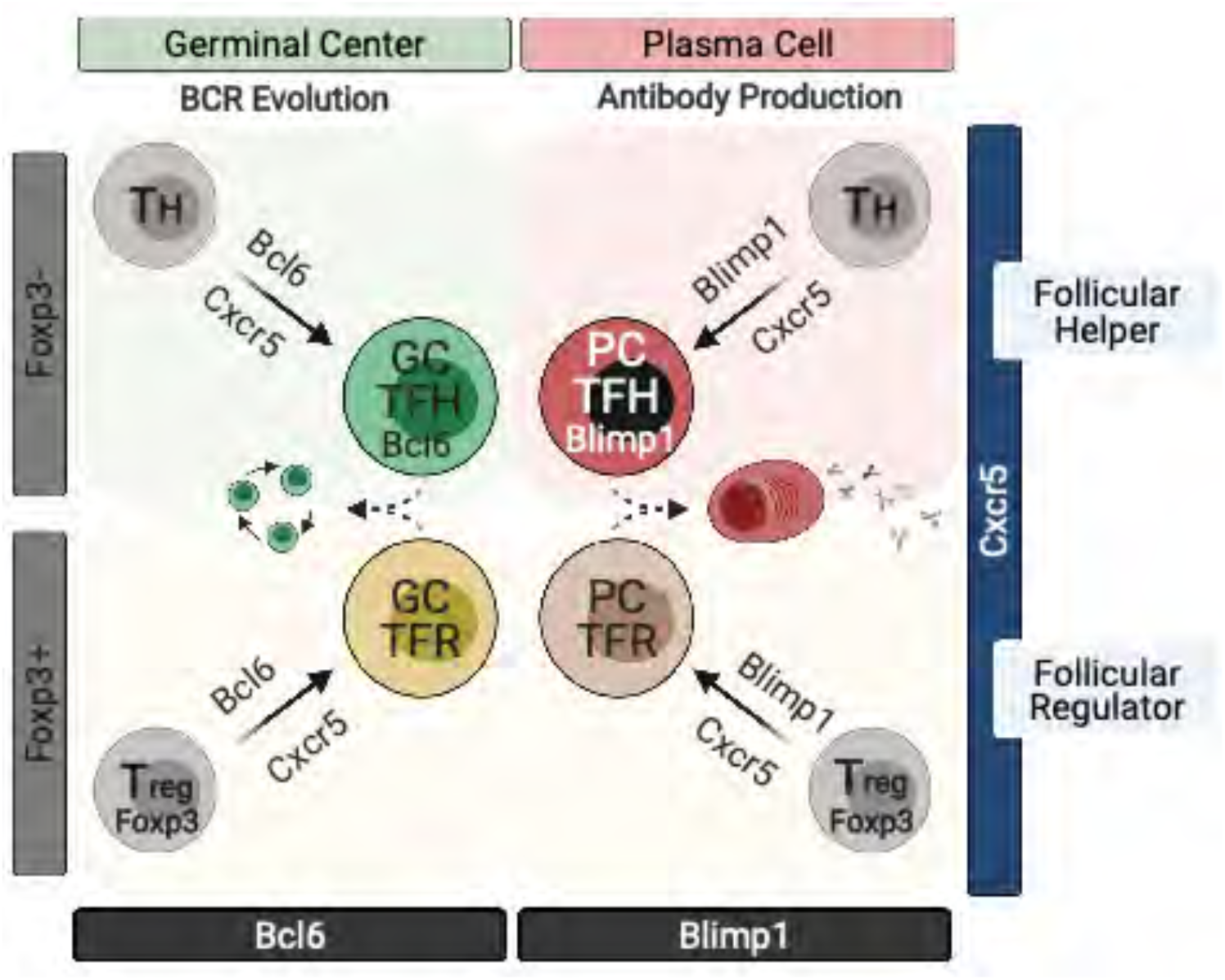

---

Circulating antibodies that have been selected and redesigned to target previous exposure to foreign antigens are the major determinants of long-term adaptive immune protection (*1–3*). This highly regulated outcome of adaptive B cell immunity is orchestrated by sub-specialized cohorts of follicular-localized CD4 T cells (*4–6*). Differentiation into either antibody-secreting plasma cells (PC) or germinal center (GC) B cells requires antigen-specific contact with follicular helper T (T_FH_) cells (*7, 8*). This dichotomous B cell fate decision requires B cell intrinsic expression of reciprocally antagonistic transcriptional repressors Blimp1 (encoded by *Prdm1*) and Bcl6 respectively. It remains unclear what T_FH_ cell mechanisms differentially impact these dichotomous antigen-driven B cell pathways.

T cell intrinsic expression of Bcl6 is a central defining component of the GC-inducing capacity of T_FH_ cells (*9–11*). In the absence of Bcl6, there is no GC formation, no BCR affinity maturation and no production of high-affinity memory B cells or post-GC PC (*4, 12*). However, it was apparent, even from the earliest germline ablation studies, Bcl6 was not required for PC differentiation (*13–15*). Across multiple germline KO models there was exaggerated type 2 helper T cell expansion and elevated PC formation in the complete absence of Bcl6. In T cell conditional deletion of Bcl6 with either adoptive transfer or within intact models, there was evidence for early residual (*16, 17*) and in some cases substantial (*18, 19*) continued support for T_H_ dependent antibody production. Recent studies of human infection demonstrate neutralizing antibody production even with diminished T_FH_ cells and GC formation (*20–22*). There is also evidence for separable cellular (*23*) and molecular pathways (*8, 24*) of T_H_ cells controlling the GC to PC fate decision. Nevertheless, Bcl6 expressing T_FH_ cells are needed for GC formation and all downstream functions of the GC cycle, however we propose a separable transcriptional program drives PC differentiation.

Blimp1 is highly expressed in CD4 T cells and is associated with both effector and regulatory T cell functions. T cell conditional deletion of *Prdm1* dysregulates immune homeostasis (*25, 26*), decreases IL-10 production in T_reg_ and T_H_1 cells and dampens the inflammatory program of T_H_17 cells. In the context of infection, Blimp1 promotes terminal differentiation in CD8 T cells (*27*), assorts with decreased IL-2 production and the non-T_FH_ CD4 T cell response that traffics to sites of antigen entry (*28*). Blimp1 is co-expressed with Bcl6 in Foxp3^+^ follicular regulatory T (T_FR_) cells (*29, 30*) that regulate T_FH_ expansion, impact antibody-mediated autoimmunity, repress early PC formation and shape GC B cell clonal evolution (*31, 32*). Recent studies demonstrate T_FR_ compartments can repress PC formation within the GC and regulate post-GC autoantibody development (*33*). In addition, T_FR_ can express the immunizing antigen specificity (*34*) and Foxp3 expressing GC-T_FR_ cells can emerge from Foxp3^-^ GC-T_FH_ to regulate GC shutdown at the terminal phase of the primary GC response (*35*). We propose a separate regulatory role for Blimp1 in T_FR_ cells and hypothesize a central division of function within the T_FH_ pathway where Blimp1 expression directs the induction of antigen-specific PC.

Here, we provide multiple layers of evidence for the bifurcation of a Blimp1 expressing PC-T_FH_ pathway of differentiation with shared and divergent elements of a transcriptional program that segregates from Bcl6^+^ GC-T_FH_ cells. In steady-state spleens, we observe Blimp1^+^ Foxp3 expressing T_FR_ cells as a putative PC-T_FR_ compartment with a related GITR^+^Foxp3^-^ PC-T_FH_ subset expressing intermediate regulatory transcriptional signatures. Primary and recall immunization induces follicular-localized Blimp1^+^Bcl6^-^ PC-T_FH_ compartments with antigen-specific PC-T_FH_ cells that rapidly emerge and transcriptionally segregate from Blimp1^-^Bcl6^+^ GC-T_FH_ cells. Therefore, persistent Blimp1 expression within Cxcr5^+^PD1^hi^ T_FH_ compartments programs the development of a separate sub-class of follicular CD4 T cells specialized to induce antigen-specific PC differentiation.

## Blimp1 expressing CXCR5^+^PD1^hi^ T_FH_ compartment

Steady-state murine spleens contain small but demonstrable levels of class-switched IgG B cells assorted across GC, memory B and PC compartments that emerge under the control of multiple follicular CD4 T cell compartments. Direct intracellular staining for Foxp3 protein and the activation marker CD44 permits separation of Foxp3^+^ CD4 T cell regulators from Foxp3^-^ CD44^hi^ TCRβ expressing non-naive effector/memory T cell compartments (**Fig. 1A**; referred to here as effector T_H_; ETH). CXCR5, as the cardinal indicator of both splenic T_FH_ and T_FR_ cells, labels similar but lower levels of these follicular T cell subsets than CXCR5^-^ ETH and regulatory T (T_reg_) compartments at steady state. Direct intracellular labeling for Blimp1 protein, reveals significant steady-state Blimp1^+^ fractions within all splenic CD4 T cell compartments (**Fig. 1A**). Although Blimp1^+^ ETH comprise the largest splenic compartment, only a small fraction of all steady-state splenic ETH expresses this transcriptional repressor. Similarly, Foxp3^+^ regulatory CD4 T cells co-express Blimp1 with distribution between CD25^hi^ and CD25^lo^ T_reg_ and T_FR_ compartments (**Fig. S1A**). Notably, Blimp1 was expressed on separable fractions of CD25^lo^ CXCR5^+^ splenic T_FH_ compartments not expected to co-express Bcl6.

**Figure 1.**
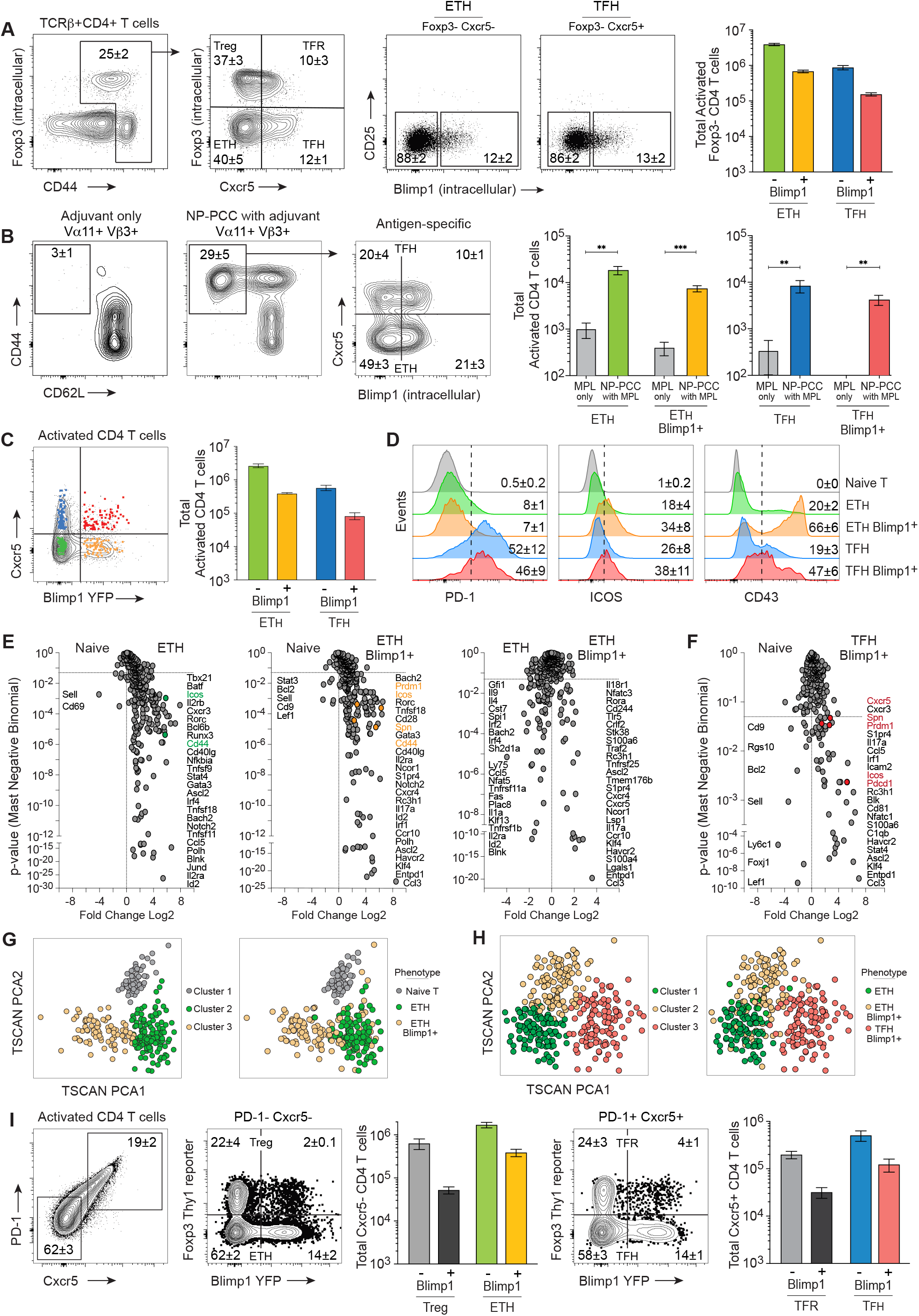
Blimp1 expressing T_FH_ cells segregate from non-T_FH_ effector cells. **A)** Flow cytometry analysis of Blimp1 expressing E_TH_ (Cxcr5^-^) or T_FH_ (Cxcr5^+^) compartments within a steady state spleen using intracellular staining. **B)** B10.BR mice were immunized with 400μg NP-PCC in MPL adjuvant or MPL alone and sacrificed on day 5. Antigen-specific Vα11^+^Vβ3^+^ CD4 T cells from the draining lymph nodes were analyzed by flow cytometry for expression of Blimp1 (intracellular) and Cxcr5 and total numbers of each subset quantified. **C)** Overlay of index-sorted splenocytes of a Blimp1-YFP reporter mouse sorted by flow cytometry for single cell qtSEQ analysis of Cxcr5^-^ E_TH_ (green or yellow) or Cxcr5^+^ T_FH_ (blue or red) subsets and **D)** protein expression flow cytometry histograms from total populations from (C). **E)** Volcano plot display of differential gene expression of E_TH_ subsets or **F)** T_FH_ subsets from single cells analyzed in (C). **G**) TSCAN pseudotime clustering of single cell RNAseq (qtSEQ) generated transcriptional programming of E_TH_ relative to naïve or **H)** Blimp1^+^ T_FH_. **I)** Splenocytes of unimmunized dual Blimp1-YFP/Foxp3-Thy1.1 reporter mice analyzed through flow cytometry displaying Blimp1-YFP protein expression within Cxcr5^-^PD-1^-^ (E_TH_ or T_reg_) and Cxcr5^+^PD-1^+^ (T_FH_ or T_FR_) compartments. Student’s *t* test: **P <0.01, ***P <0.001. Flow plots display mean±SEM. Data are representative of 2-3 independent experiments with n=3-5 total mice.

Using a well-established protein immunization model (*36, 37*), we reveal substantial fractions of antigen-reactive Blimp1 expressing ETH as early as day 5 after initial priming (**Fig. 1B**). These ETH compartments rapidly exit the draining LNs and without CD62L are unable to re-enter. At this early stage of the immune response, substantial fractions of both Blimp1^-^ and Blimp1^+^ T_FH_ cells accompany rapid and early antigen-specific PC differentiation prior to GC formation (**Fig.1B**). Both T_FH_ compartments continue to expand by day 7 of the primary response but remain at similar 2:1 ratios (**Fig. S1B**). Hence, while Blimp1 expressing ETH comprise substantial fractions of the response to protein antigen, there were also emergent antigen-reactive Blimp1 expressing CXCR5^+^ T_FH_ compartments.

To confirm and extend analysis of the steady-state Blimp1^+^ T_FH_ compartment, we use a Blimp1-YFP reporter strain. In this strain, we use surface expression of TNFR-related protein GITR, that correlates positively with Foxp3 expression, to separate GITR^-^ T_FH_ from GITR^+^ T_FR_ splenic compartments. Using reporter expression, we find similar numbers and fractions of Blimp1^-^ and Blimp1^+^ compartments of ETH and T_FH_ as observed using direct intracellular staining for protein (**Fig. 1C**). Using this model, we demonstrate that PD1 expression is differentially expressed on both T_FH_ subsets (**Fig. 1D**) indicating that these CXCR5^+^ compartments are most likely within GC microenvironments (*38*). All non-naive T cell compartments express higher levels of ICOS while the Blimp1^+^ ETH compartment contained a large subset with distinctively high levels of CD43. Thus, persistent Blimp1 expression in the steady-state spleen separates a subset of the ETH compartment and further segregates a Blimp1^+^ CXCR5^+^PD1^hi^ T_FH_ cell compartment.

## Separable Blimp1^+^ ETH and T_FH_ transcriptional programs

Using the Blimp1-YFP reporter and GITR expression, we sorted Blimp1^+^ and Blimp1^-^ cells from both ETH and T_FH_ subsets. We recently developed a single cell-indexed strategy to assess gene expression within these very rare subsets of CD4 T cells at steady-state (*39*). In this strategy, we couple fluorescence-based flow cytometry to quantitative and targeted RNA sequencing (single cell qtSEQ) to directly integrate cell surface phenotype with expression of target genes. The targeted library formation uses semi-nested-PCR for high sensitivity amplification and library formation with digital deconvolution to track the cellular origin of the mRNA and for amplification adjusted quantification. Library size using this method is comparable to that seen with more global techniques, however, as these studies target ∼500 genes, the dropout rate for individual cells is lower and sensitivity is ∼10-fold higher for each assay at the single cell level. While throughput is low, single cell qtSEQ provides greater depth for the targeted assessment of transcriptional programs in rare, phenotypically-complex cellular compartments.

In this initial analysis, differential transcriptional programs segregate Blimp1^-^ and Blimp1^+^ ETH from each other and from Blimp1^+^ T_FH_ compartments at steady-state (**Fig. 1E**). As expected, major transcriptional regulators of known effector CD4 T cell programs such as *Tbx21*, *Batf*, *Rorc*, *Gata3* are readily detectable in the Blimp1^-^ ETH compartment. Other major transcriptional regulators with known inducible (eg *Bach2, Stat4, Irf4, Ascl2*) or negative impact on lymphocyte differentiation (eg *Bcl6b, Id2, Runx3*), expression of multiple surface determinants of function (eg *Icos, Cd44, Cxcr3, Il2ra, Il2rb*) and molecular co-ordinators of cell-cell interactions (eg *Tnfsf9*, *Tnfsf18*, *Tnfsf11*) are also elevated in expression in the Blimp1^-^ ETH compartment. Differential analysis highlights increased levels of transcriptional regulators in Blimp1^+^ ETH (eg *Rora, Ascl2, Ncor1, Klf4*) over the ETH (eg *Spi1, Id2, Klf13, Irf4, Irf2, Bach2*). Expression for surface molecules emphasizes distinctions in Blimp1^+^ ETH for a range of chemokine receptors (eg *S1pr4, Cxcr4, Cxcr5, Ccr10*) and immune function regulators (eg *Havcr2*, Tim3; *Tnfrsf25*, DR3; *Tlr5*) over the Blimp1^-^ ETH compartment (eg *Sh2d1a*, *Ly75, Fas, Il2ra, Tnfsf11a*, *Tnfrsf1b*). Using principal component analysis with gene expression alone, sorted single cells from Blimp1^-^ and Blimp1^+^ ETH segregate from naive T_H_ cells and from each other (**Fig. 1G**), further highlighting the divergent nature of CD4 non-follicular effector/memory subsets in the steady-state spleen.

Blimp1^+^ T_FH_ compartments also segregated from naïve T and both ETH subsets (**Fig. 1F**). Contributing to these differences are notable transcriptional regulators (eg *Prdm1, Klf4, Ascl2, Stat4, Nfatc1* and *Irf1*, surface modulators of location (eg *Cxcr5, Cxcr3, S1pr4*) and intercellular contact (eg *Pdcd1, Havcr2, Spn*, *Cd81*, *Icam2, Entpd1*) with differential expression of cytosolic machinery (eg *Rc3h1*, *S100a6*, *Blk*) and secreted molecules (eg *Ccl3, Ccl5*) prominent in the Blimp1^+^ T_FH_ compartment. These trends introduce broad patterns of divergence at a level of the transcriptional program that potentially regulate separable cellular functions.

## Lymphoid assortment of Blimp1^+^ follicular T cell sub-compartments

Next, we developed a dual reporter model introducing both Blimp-YFP and Foxp3-Thy1.1 together to separate T_FH_ and T_FR_ cells and their respective Blimp1 expressing sub-compartments. As expected, there are lower numbers of conventional T_reg_ cells in female spleens, however, the ratio of Blimp1^-^ to Blimp1^+^ appears similar to males (**Fig. 1I** **& S1B**). In this dual reporter model, there were similar numbers and dominance of Blimp1^-^ CXCR5^-^PD1^-^ ETH over the Blimp1^+^ sub-compartment as seen in the single Blimp1 reporter using GITR expression to exclude T_reg_. The follicular compartments were larger overall in female spleens with similar numbers of Blimp1^-^ T_FR_ but lower fractions and number of Blimp1^+^ T_FR_ cells. There were similar compartments of Blimp1^-^ T_FH_ but consistently higher numbers and fractions of Blimp1^+^ T_FH_ cells in female spleens (**Fig. 1I**). Small but discernible fractions of Blimp1^+^ CXCR5^+^PD1^hi^ T_FH_ and T_FR_ compartments can be found consistently in the bone marrow, mesenteric LNs and Peyer’s Patch lymphoid tissues (**Fig. S1B**). While the balance altered considerably, there was consistent presence of both Blimp1 expressing T_FH_ and T_FR_ compartments across central and peripheral lymphoid tissues at homeostasis.

## CD4 T cell expressed Prdm1 is required for PC differentiation

*Prdm1* is a major transcriptional repressor central to the formation and function of antigen-specific PC differentiation in B cells (*40*). As described previously, germline conditional ablation of Prdm1 in T cells introduces complex interdependent phenotypes with multiple levels of impact at T cell development, peripheral regulatory and effector T_H_ cell functions (*25, 26*). Here, we chose an adoptive transfer model in which naive T_H_ cells transferred into immunodeficient RAG1^def^ recipients become activated, expand and differentiate into a spectrum of B cell helper compartments. Resultant CD4 T cells drive class-switched PC differentiation within 14 days and delayed GC formation occurs by day 28 after transfer with or without intentional immunization (*23*). Here, we sorted and separated CD4 T cells and CD19^+^ B cells from Prdm1^fl/fl^-Rosa26^CreERT2^ spleens. This mature CD4 T cell compartment was treated with tamoxifen or vehicle in vitro for 1h driving >90% deletion of target gene, before adoptive co-transfer with untreated splenic B cells.

In the first series of studies, we targeted a late timepoint to permit GC formation and sufficient time to decrease short-lived PC known to have 3-5 day half-life in vivo . At day 28, the reactive CD4 T cell compartment was >90% activated CD44^hi^CD62L^lo^ T cells with similar numbers regardless of treatment (**Fig. 2A**). Distribution of CD25 and CXCR5 was also broadly similar suggesting the capacity for a broad range of functional outcomes (**Fig. 2B**). The transferred T cell compartments supported equivalent amounts of GC formation (**Fig. 2C**), no change in memory B cell numbers (**Fig. S2C**) but significantly truncated class-switched PC differentiation (**Fig. 2D**). Next, we targeted the delayed pre-GC PC differentiation pathway at day 14 after treatment and transfer. Transferred splenic B cells, expand with predominantly IgD^-^IgM^+^ B cells and IgD^-^IgM^-^ class-switched B cells present (**Fig. S2D**) before any GL7^hi^CD38^lo^ GC formation (**Fig. S2E**). We detected substantial baseline PC formation that is likely T cell independent after transfer into these immunodeficient recipients. Nevertheless, there was significant truncation of both IgM and class-switched PC formation after CD4 conditional and tamoxifen-induced temporal deletion of *Prdm1* (**Fig. 2E,F**). This flow cytometry-based analysis was confirmed using IgG specific EliSPOT assessment for frequencies of antibody-secreting cells (**Fig. 2G**). Taken together, these studies provide a causal link between CD4 T cell expression of *Prdm1* and both pre-GC and post-GC pathways of PC differentiation in vivo.

**Figure 2.**
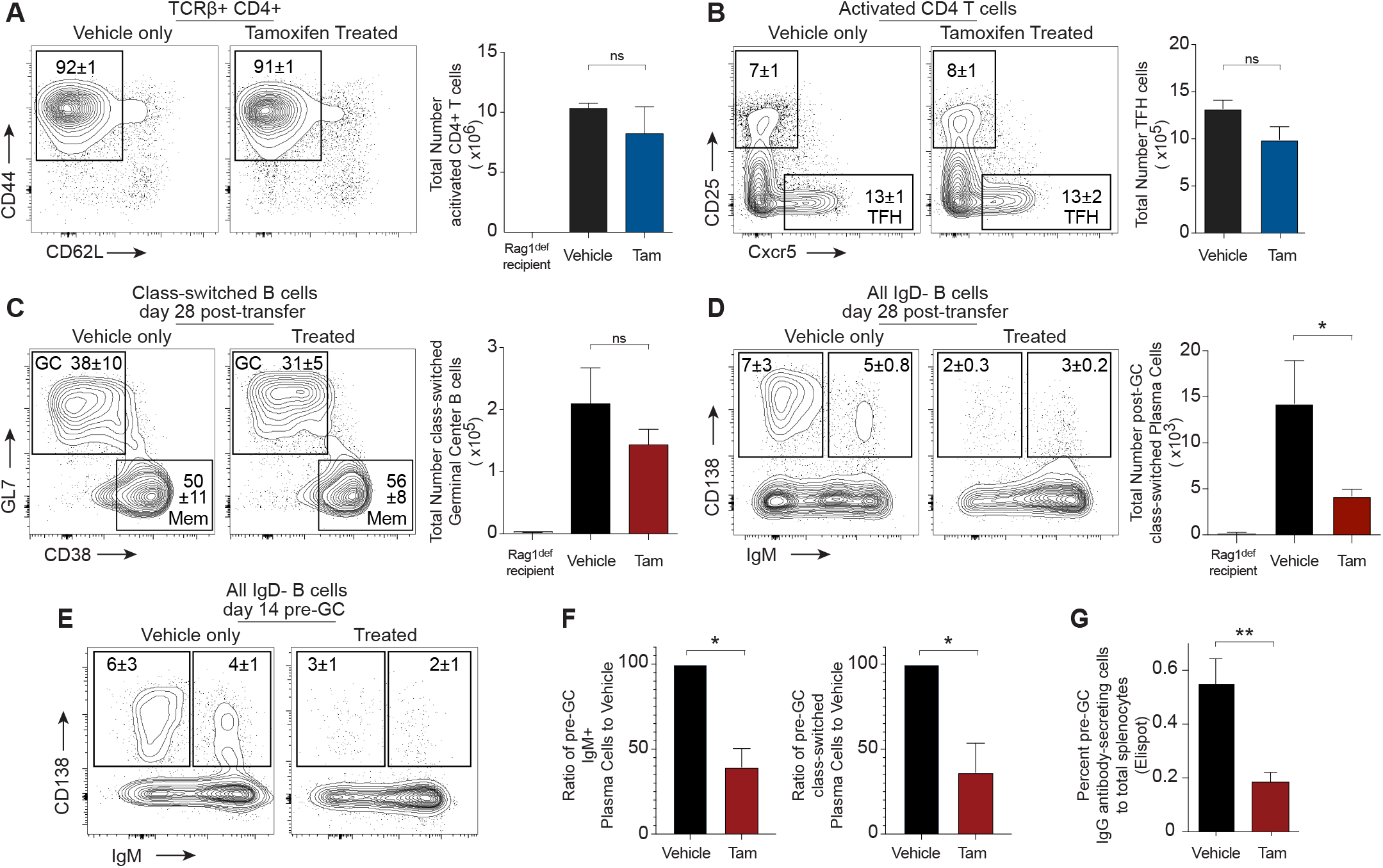
CD4 T cell expressed *Prdm1* is required for PC differentiation in vivo. Adoptive transfer of Prdm1^fl/fl^ CD4^+^ T splenocytes treated with or without tamoxifen (2.5x10^6^) and untreated B cells (1x10^7^) transferred i.p. into Rag1^def^ recipients and immunized with 400μg NP-KLH in adjuvant. Mice were sacrificed and analyzed on day 14 or 28. **A)** Analysis of total TCRb^+^CD4^+^ T cells and **B)** activated CD44^+^ T cell expression of CD25 or Cxcr5 at day 28, depicted by bar graphs of total numbers and flow cytometry plots. **C)** Separation of GC (CD19^+^B220^+^GL7^+^CD38^-^) or memory (CD19^+^B220^+^GL7^-^CD38^+^) subsets of class-switched (IgM^-^ IgD^-^) B cells and **D)** total class-switched or IgM^+^ plasma cells (CD138^+^) from day 28 post-transfer of vehicle or tamoxifen-treated splenocytes depicted as representative flow plots (left) or bar graphs (right). **E)** Analysis of class-switched (IgD^-^IgM^-^) and IgM^+^ (IgD^-^IgM^+^) plasma cells from day 14 post-transfer displayed as flow cytometry plots, **F)** relative expression of tamoxifen-treated cell numbers to vehicle and **G)** secreted IgG by Elispot from the spleen. Student’s *t* test: ns = not significant, *P <0.05, **P <0.01. Flow plots display mean±SEM. Data are representative of three independent experiments with n= 6-7 total mice per timepoint.

## CD25^lo^Blimp1^+^ splenic CD4 T cells promote PC differentiation

To narrow down which steady-state mature splenic CD4 T cell compartment was responsible for PC differentiation, we established in vitro activated T_H_-B cell co-cultures (**Fig. 3A**). Sorted T cells are polyclonally activated in vitro for 2 days and then ’rested’ without TCR triggers in IL2 alone for 2 days before addition of Blimp1^-^ CD19^+^ B cells for a further 5 days prior to assay for PC differentiation. Using the Blimp1-YFP reporter spleens, we isolated GITR^-^ Blimp1^-^ CD4 T cells (ETH and conventional T_FH_), Blimp1^+^GITR^-^ CD4 T cells (ETH and T_FH_) and Blimp1^+^ GITR^+^ CD4 T cells (Blimp1^+^ T_reg_ and T_FR_)(**Fig. 3B**). Naive T cells were used in all co-cultures and help to establish baseline PC differentiation without addition of non-naive T cell subsets. At day 9, in vitro co-cultured T cells were assessed for Blimp1-YFP expression by flow cytometry (**Fig. 3C**). Some wells defined as Blimp1^-^ T cells at the time of sorting upregulated Blimp1 in vitro (∼64%), However, no naive T cells upregulated Blimp1 in these cultures and most Blimp1 expression correlated with pre-existing levels on sorted populations.

**Figure 3.**
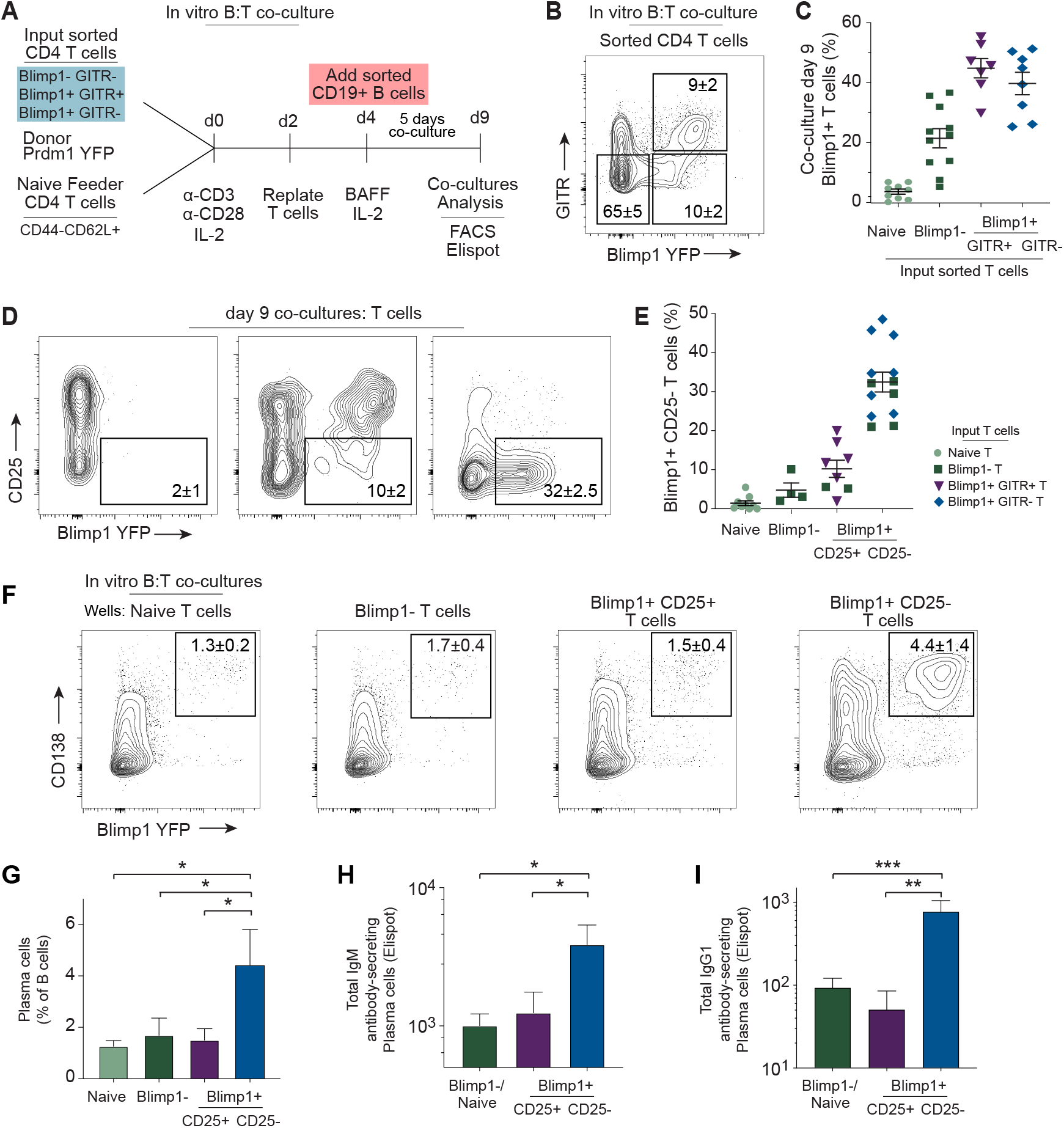
Blimp1^+^CD25^lo^ CD4 T cells promote highest PC differentiation in vitro. Blimp1^-^, Blimp1^+^GITR^+^, Blimp1^+^GITR^-^ and naïve TH subsets were sorted and cultured with CD3, CD28, and IL-2 for 48 hours, rested for 48 hours, and then combined with Blimp1^-^CD19^+^ B cells in the presence of BAFF and IL-2 for 5 days to generate T-B co-cultures. **A)** Experimental design of T-B cell cocultures. **B)** T-cell sorting strategy of distinct Blimp1/GITR subset phenotypes depicted as a representative flow cytometry plot and **C)** levels of Blimp1 expression in each subset, organized by original sort population (B). Subsets are arranged by shape and color: green circle, naïve T; green square, Blimp1^-^TH; purple triangle, Blimp1^+^GITR^+^TH; blue diamond, Blimp1^+^GITR^-^ TH. **D)** Analysis of co-cultured T cells re-ordered by phenotypic Blimp1 expression at day 5 of co-culture (day 9 culture) depicted as flow cytometry plots displaying Blimp1 and CD25 expression and **E)** dot plot display of Blimp1^+^CD25^-^ T cells re-ordered into Naive, Blimp1^-^, Blimp1^+^CD25^+^, or Blimp1^+^CD25^-^ subsets by Day 5 phenotype but depicted as symbols of original sort described in (B, C). **F)** Multi-dimensional display of plasma cell differentiation from day 5 co-culture subsets from (D), gated as Blimp1^+^CD138^+^ through flow cytometry, **G)** summarized in a bar graph as a percentage of total B cells, **H)** IgM and **I)** IgG1 secretion from Elispot assays. Student’s *t* test: ns = not significant, *P <0.05, **P <0.01. Flow plots display mean±SEM. Data are representative of three independent experiments with across 32 total wells.

The CD4 T cells with newly-expressed Blimp1 assorted as either CD25^+^ or CD25^lo^ with the latter representing highest penetrance across all conditions (**Fig. 3D,E**). Using both CD138 and Blimp1 expression on IgM and class-switched B cells as a measure of PC differentiation, we found significant amounts of IgM, non-IgM class-switched and IgG1 PC only in co-cultures containing in vitro activated CD25^-^Blimp1^+^ CD4 T cells (**Fig. 3F,G**). In these studies, IgM and IgG1 secretion was confirmed using specific EliSPOT assays (**Fig. 3H**). Thus, in vitro T cell functional assays correlate Blimp1 expression in CD25^lo^ T_H_ cells to the promotion of both IgM^+^ and class-switched PC differentiation.

## Separable PC-T_FH_ and GC-T_FH_ transcriptional programs

Using the dual reporter model, we separated Foxp3^-^ and CXCR5^+^PD1^hi^ T_FH_ compartments from steady-state spleens based on Blimp1-YFP expression for single cell qtSEQ analysis (**Fig. 4A**). These high-fidelity single cell sorts selected CD4^+^TCRβ^+^ T cells expressing high levels of CD44 and PD1 that were mostly ICOS high, with varying levels of CD62L that largely correlated with Ly6C (**Fig. S3A**). Foxp3^-^Blimp1^-^ T_FH_ cells were all GITR^-^ CD25^-^ and will be referred to here as the GC-T_FH_ compartment. In contrast, Foxp3^-^Blimp1^+^ T_FH_ cells were further divided by GITR and CD25 expression with the GITR^-^CD25^lo^ compartment most closely aligning with PC-inducing function and will be referred to here as the PC-T_FH_ compartment. TSCAN analysis using gene expression only and PCA broadly segregates the single cell transcriptional programs of GC-T_FH_ and PC-T_FH_ from naive T cells and each other (**Fig. S3B**), while volcano plots highlight significant differences between the two major T_FH_ subsets (**Fig. S3C**). However, separate comparisons of each T_FH_ subset to naive T cells reveals multiple shared and many uniquely upregulated elements that define and differentiate the GC-T_FH_ and PC-T_FH_ transcriptome at homeostasis (**Fig. 4B, S3C-F**).

**Figure 4.**
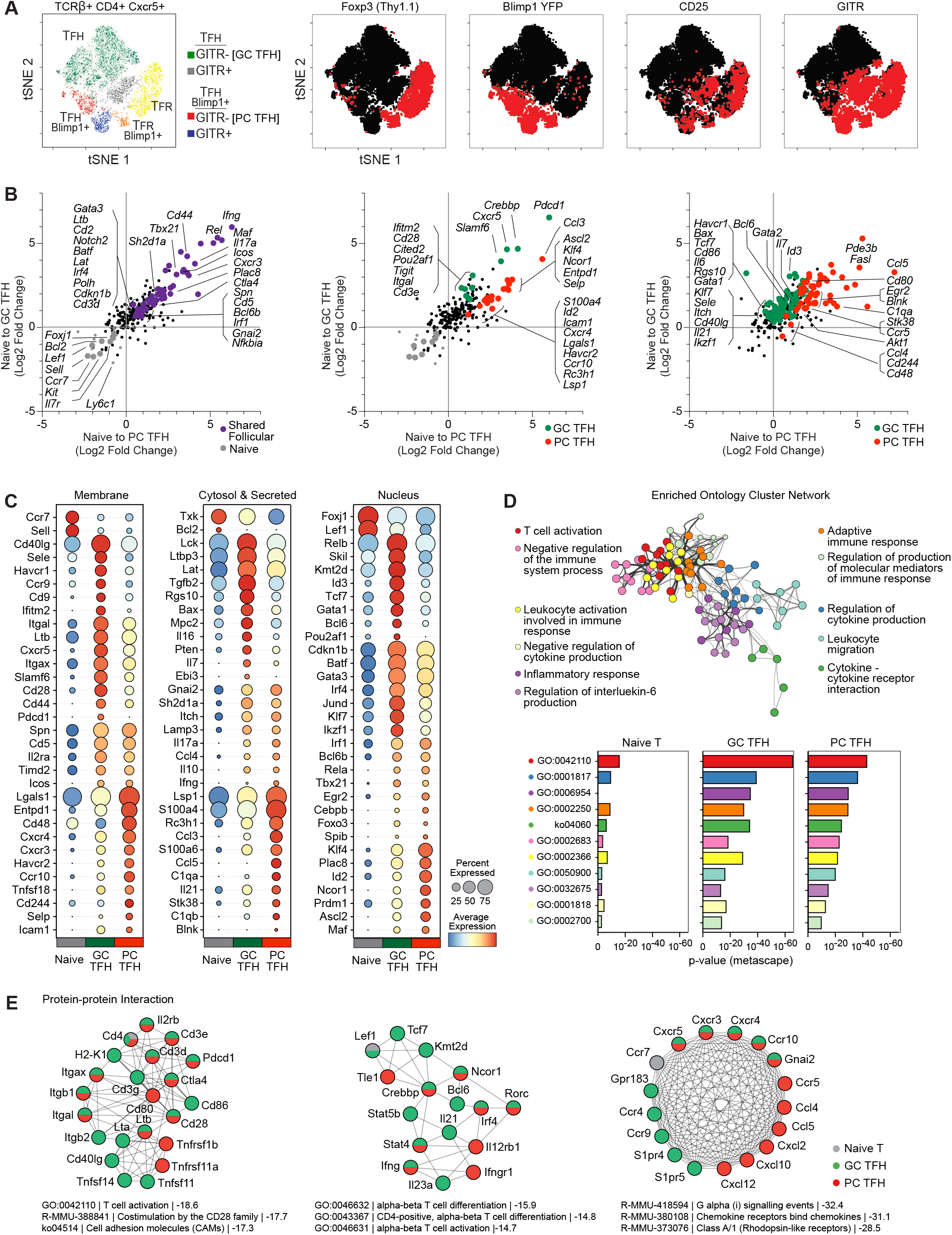
PC T_FH_ and GC T_FH_ maintain separable transcriptional programs at steady-state. **A)** tSNE dimensionality reduction of Cxcr5^+^ T_FH_ splenocytes from unimmunized dual Blimp1-YFP/Foxp3-Thy1.1 reporter mice. Colors indicate phenotype: green, Blimp1^-^T_FH_ (GITR^-^); grey, Blimp1^-^T_FH_ (GITR^+^); yellow, T_FR_ (Blimp1^-^); orange, T_FR_ (Blimp1^+^); blue Blimp1^+^T_FH_ (GITR^+^); red Blimp1^+^T_FH_ (GITR^-^). tSNE display is separated by phenotype (left) and overlayed individual protein expression of Foxp3, Blimp1, CD25 or GITR (right). **B)** Scatter plot of single cell RNA-seq (qtSEQ) differential gene expression in naïve to PC-T_FH_ (x-axis) and naïve to GC-T_FH_ (y-axis). Colored dots indicate p-value <0.05. Plots display commonly upregulated genes with no significant difference between GC-T_FH_ and PC-T_FH_ expression (left; blue), commonly upregulated genes, but significantly different GC-T_FH_ and PC-T_FH_ expression (center; green, GC-T_FH_; red, PC-T_FH_), and uniquely upregulated genes in either GC-T_FH_ or PC-T_FH_ (right; green, GC-T_FH_; red, PC-T_FH_). **C)** Dot plot display of average gene expression within naïve T, GC-T_FH_, and PC-T_FH_ subsets. Dot color reflects average gene expression and dot size depicts population penetrance. **D)** Metascape visualization of the interactome network of commonly upregulated GC and PC T_FH_ pathways and expression of GC-T_FH_, PC-T_FH_ or naïve within each Gene Ontology (GO) pathway or **E)** protein-protein interaction network expression. The size of each node positively correlates with the number of edges it connects to. Clusters are colored according to their enriched gene ontology pathway or functional identities (green, GC-T_FH_; red, PC-T_FH_; grey, Naïve T). The molecular complex detection (MCODE) algorithm identifies individual genes that are densely associated with a larger functional unit. Dot size correlates to Log2 fold change expression of GC-T_FH_ or PC-T_FH_ relative to naïve. Single cell RNAseq analysis (qtSEQ; B-E) included post-QC subsets of GC-T_FH_ (n=304), PC-T_FH_ (n=248) and naïve T (n=153) from two independent experiments with 3-4 mice per experiment.

To display the similarities and differences between GC-T_FH_ and PC-T_FH_ we used circle plots that represent not only average gene expression but also percent positive for targeted assays (**Fig 4C, S3G**). Results have been ordered hierarchically to emphasize gene targets with highest expression and greatest penetrance and separated broadly by where the protein product is expressed within the cell (membrane, cytosol/secreted and nuclear). As expected, multiple mRNA species expressed in naive T cells significantly decrease upon T_FH_ differentiation into either pathway (eg *Ccr7, Sell, Txk, Bcl2, Foxj1, Lef1*). Many target genes were significantly more elevated within GC-T_FH_ and serve to emphasize GC-inducing T_FH_ functions. Membrane expressed molecules involved in co-stimulation (eg *Cd40lg, Cd28, SlamF6*), cell-cell adhesion (eg *Cd44, Cd9*, *Itgax, Itgal, Itgb2*), guidance cues (eg *Cxcr5, Ccr9, Gpr183*) and known modifiers of cell fate (eg *Ltb, Havcr1*, *Tigit, Ptprc, Cd27, Tnfrsf8*) were higher in GC-T_FH_.

Surface molecules and intracellular machinery associated with TCR activation increases in GC-T_FH_ (eg *Cd3ε, Cd3γ, Cd3δ, Lat, Lck*) may indicate greater dynamic activity within splenic GC than induction of PC differentiation at homeostasis. At the level of transcriptional regulators, GC-T_FH_ cells expressed significantly higher amounts of many known factors previously associated with the T_FH_ pathway (eg *Bcl6, Tcf7, Tox*) and others involved in controlling T cell/B cell differentiation (eg *Relb, Id3, Pou2af1, Gata1*) with elevated expression of genes associated with effector/memory lymphocyte differentiation (eg *Batf, Gata3, Irf4, Ikzf1*). These increased cellular activities reflect the status of splenic GC with ongoing adaptive tolerance mechanisms and acute immune effector functions, even at steady-state.

Shared expression of surface (eg *Spn, Cd5, Il2ra, Icos*) and secreted/cytosolic components (eg *Gnai2, Sh2d1a, Itch, Ccl4, Il10*) emphasize potential for common T cell physiology and B cell helper functions. Similar levels of known lymphocyte regulators (eg *Tbx21, Irf1, Bcl6b, Rela, Egr2*) also indicate shared persistent functions between cells in these pathways at steady-state. However, differential expression of cell surface, secreted and cytosolic components of the PC-T_FH_ pathway can also be detected. PC-T_FH_ cells express higher levels of different costimulatory molecules (eg *Cd48, Entpd1, Cd244*), adhesion machinery and guidance receptors (eg *Icam1, Cxcr3, Cxcr4, Ccr10*) and known modifiers of cell fate (eg *Ctla4, Tnfsf18, Havcr2*) than GC-T_FH_. Some secreted factors (eg *Ccl5, Ccl6, Il21, Tnfsf13*) and cytosolic elements involved in survival and migration (eg *Rc3h1, Lsp1*) were also differentially expressed in the PC-T_FH_ compartment. Regarding transcriptional regulators, as expected PC-T_FH_ differentially expressed *Prdm1* with other known major modifiers of lymphocyte programming (eg *Ascl2, Klf4, Id2, Ncor, Maf, Plac8*). Hence, shared and unique elements of the PC-T_FH_ program begin to elucidate differential follicular mechanisms controlling T_FH_ cell dependent PC differentiation.

To emphasize the shared attributes of these follicular T cell compartments, network analysis using Metascape (*41*) revealed multiple enriched onotology clusters involving T cell activation, positive and negative immune response mediation, and cytokine and chemokine production that are similarly significant in both pathways (**Fig. 4D**). Interrogation of predicted protein-protein interactions (**Fig. 4E**) revealed common pathways used by both subsets involving T cell activation pathways, co-stimulation, CD4 T cell differentiation and chemokine and cytokine regulation. However, while many of the molecular components in these pathways were shared between subsets, multiple others were uniquely expressed. These trends indicate mechanistic differences in how each compartment is reacting to, or has previously reacted to, immune challenge. Taken together, these results indicate multiple shared and divergent elements of follicular T cell transcriptional programing that reveal the bifurcation of GC-T_FH_ and PC-T_FH_ pathways of differentiation.

## Segregated Blimp1 expressing T_FR_ regulatory program

As discussed, Blimp1 expression is most typically associated with Foxp3 expressing T_FR_ cells that exert a negative regulatory function on B cell differentiation (*29, 30*). Using the dual reporter model, we could readily segregate a minor Blimp1^+^ T_FR_ subset from the majority Blimp1^-^ T_FR_ compartment (**Fig. 4A**, **S4A**). However, there was also a substantial compartment of Foxp3^-^ Blimp1^+^ T_FH_ cells that expressed GITR, a protein also associated with regulatory function (**Fig. S4A**) . Based on differential protein expression, including higher Blimp1, PD1, CD25, ICOS and lower Ly6C, these GITR^+^Blimp1^+^ T_FH_ cells most closely track with the Blimp1^+^ T_FR_ compartment and predict regulatory function (**Fig. S4B**). In support of this division, the GITR^-^ PC-T_FH_ express higher levels of T_FH_ associated transcriptional regulators *Bcl6* and *Tcf7* while the GITR^+^Blimp1^+^ T_FH_ subset express multiple immune checkpoint molecules such as *Ctla4, Tnfrsf18, Havcr2* and *Tigit* (**Fig. S4C**). More broadly, the transcriptional program of these cells, referred to here as GITR^+^ PC-T_FH_ cells, express more distinct elements from GC-T_FH_ and PC-T_FH_ compartments and greater overlap with the Blimp1^+^ T_FR_ compartment (eg *Tnfsf18, Ctla4, Havcr2, Ikzf4, Klrg1* and *Prdm1*) (**Fig. 5A, S4D**).

**Figure 5.**
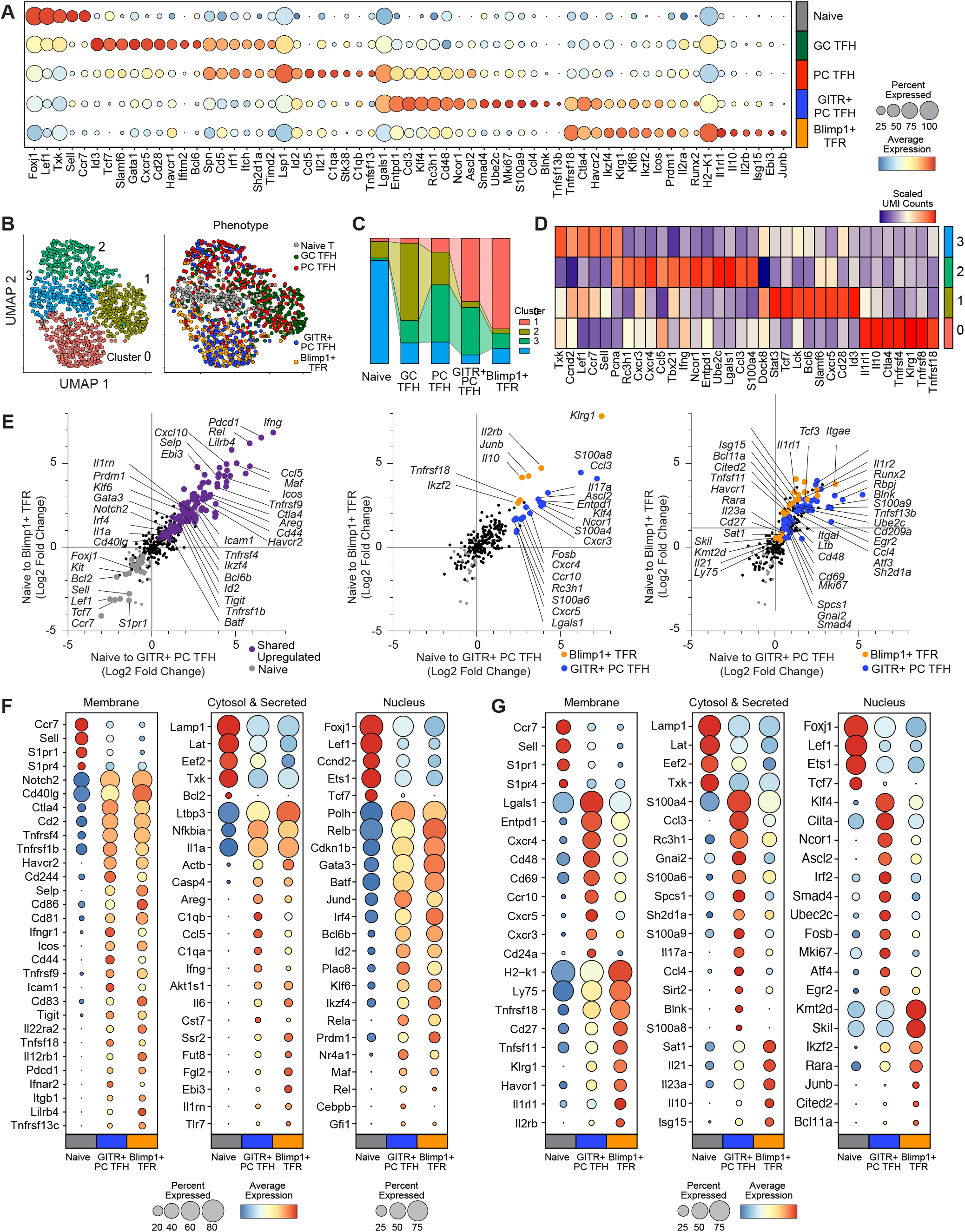
Blimp-1 expressing Foxp3^+^ putative PC T_FR_ subset. **A)** A dot plot depiction of average gene expression across all subsets analyzed using single cell RNAseq (qtSEQ) from unimmunized dual Blimp1-YFP/Foxp3-Thy1.1 reporter spleens. Dot color reflects average gene expression and dot size depicts population penetrance. **B)** UMAP dimensionality reduction of naïve (n=153), GC-T_FH_ (n=304), PC-T_FH_ (n=248), Blimp1^+^GITR^+^ T_FH_ (n=254), and Blimp1^+^T_FR_ (n=245) as UMAP generated clusters (left) or overlaid with sort phenotype (right), **C)** organized by subset to display proportion allocated to each UMAP cluster (cluster abundance) and **D)** heatmap display of key significant differentially expressed genes from each UMAP cluster. **E)** Scatter plot of single cell RNA-seq (qtSEQ) differential gene expression in naïve to GITR^+^ Blimp1^+^T_FH_ (x-axis) and naïve to Blimp1^+^T_FR_ (y-axis). Colored dots indicate p-value <0.05. Plots display commonly upregulated genes with no significant difference between GITR^+^ Blimp1^+^T_FH_ and Blimp1^+^T_FR_ expression (left; blue), commonly upregulated genes, but significantly different GITR^+^ Blimp1^+^T_FH_ and Blimp1^+^T_FR_ expression (center; orange, Blimp1^+^T_FR_; blue, GITR^+^ Blimp1^+^T_FH_), and uniquely upregulated genes in either GITR^+^ Blimp1^+^T_FH_ or Blimp1^+^T_FR_ (right; orange, Blimp1^+^T_FR_; blue, GITR^+^ Blimp1^+^T_FH_). **F)** A dot plot display of differentially expressed genes across naïve, GITR^+^ Blimp1^+^T_FH_ and Blimp1^+^T_FR_ depicting shared genes or **G)** unique genes, as described in (E). Dot color reflects average gene expression and dot size depicts population penetrance. Single cell RNAseq analysis from two independent experiments with 3-4 mice per experiment.

Clustering naive T_H_ cells with these four non-naive CD4 compartments further supported the alignment of GC-T_FH_ and PC-T_FH_ transcriptional programs (Cluster 1 higher shared levels *Dock8, Stat3, Tcf7, Bcl6, Slamf6, Cxcr5, Cd28*) and the GITR^+^ PC-T_FH_ with Blimp1^+^ T_FR_ compartment (Cluster 0, higher shared levels *Il10, Ctla4, Tnfrsf4, Klrg1, Tnfrsf8, Tnfrsf18*) **Fig. 5B-D**). A separate cluster (Cluster 2; *Cxcr3, Tbx21, Ifng*) comprises more shared type 1 activities across all compartments than attributes that divide T_FH_ and T_FR_. Hence, we conclude that the GITR^+^ PC-T_FH_ compartment expresses more regulatory than helper attributes and may represent a transitional state between the GITR^-^ PC-T_FH_ and its Blimp1^+^ T_FR_ counterpart.

For further assessment of the Blimp1 associated follicular regulatory program, we compared each subset to naive T_H_ cells to highlight the significantly different but shared or unique features of this regulatory program (**Fig. 5E; S4E,F**). Many well-described examples of this shared regulatory program (eg *Tnfrs4, Tigit, Pdcd1, Ctla4*) are highly expressed mRNA species associated with inter-cellular contact, control and secreted or cytosol expressed molecules (**Fig. 5F**). Shared transcriptional regulators also abound with known action in lymphocyte differentiation. In contrast, there are many examples of uniquely membrane, secreted/cytosol and nuclear expressed molecules that may define the transitional progress from the GITR^+^ PC-T_FH_ compartment towards the Blimp1^+^ T_FR_ program (**Fig. 5G**). Thus, both compartments express broadly regulatory transcriptional programs that may target PC differentiation in ways that broadly correspond to facets of GC-T_FR_ control of GC B cell function.

## Antigen-specific memory response PC-T_FH_ and GC-T_FH_ programs

We have provided multiple layers of evidence for separable PC-T_FH_ and GC-T_FH_ follicular T cell compartments after initial antigen priming and in the steady-state immune system. The steady-state spleen is further complicated by the existence of adaptive immune tolerance mechanisms that manage commensal and environmental antigens (*42, 43*). However, steady-state spleens would also contain effector B and T cell compartments reactive to foreign antigens with many expected to be in a persistently altered but quiescent memory state.

Next, we focus on sampling the transcriptional programs of antigen reactive memory T_FH_ compartments. We chose a simple protein antigen model, priming with chicken ovalbumin and a TLR4 agonist adjuvant (Monophosphoryl lipid A; MPL) with repeat immunization >10 weeks for recall. Using Ova-I-A^b^ tetramer labeling, we sorted single antigen-activated CD44^hi^ pMHCII^+^ CXCR5^+^ memory response T_FH_ cells 4 days after challenge for gene expression analysis (**Fig. 6A**). Immunofluorescence for Bcl6 and Blimp1 protein revealed CD4^+^ T cells expressing either transcription factor in the follicular B220^+^ B cell regions of reactive lymph nodes (**Fig. 6B**). Quantifying the distribution of these follicular subsets displays lower fractions of Blimp1^+^Bcl6^-^ PC-T_FH_ cells than Blimp1^-^Bcl6^+^ GC-T_FH_ following soluble versus adjuvant inclusion at recall. The significant increase in PC-T_FH_ cells with adjuvant correlates with earlier studies indicating the increased impact of MPL on memory PC expansion (*44*). Nevertheless, this imaging analysis ascertains the follicular localization of a Blimp1^+^Bcl6^-^ memory response PC-T_FH_ compartment.

**Figure 6.**
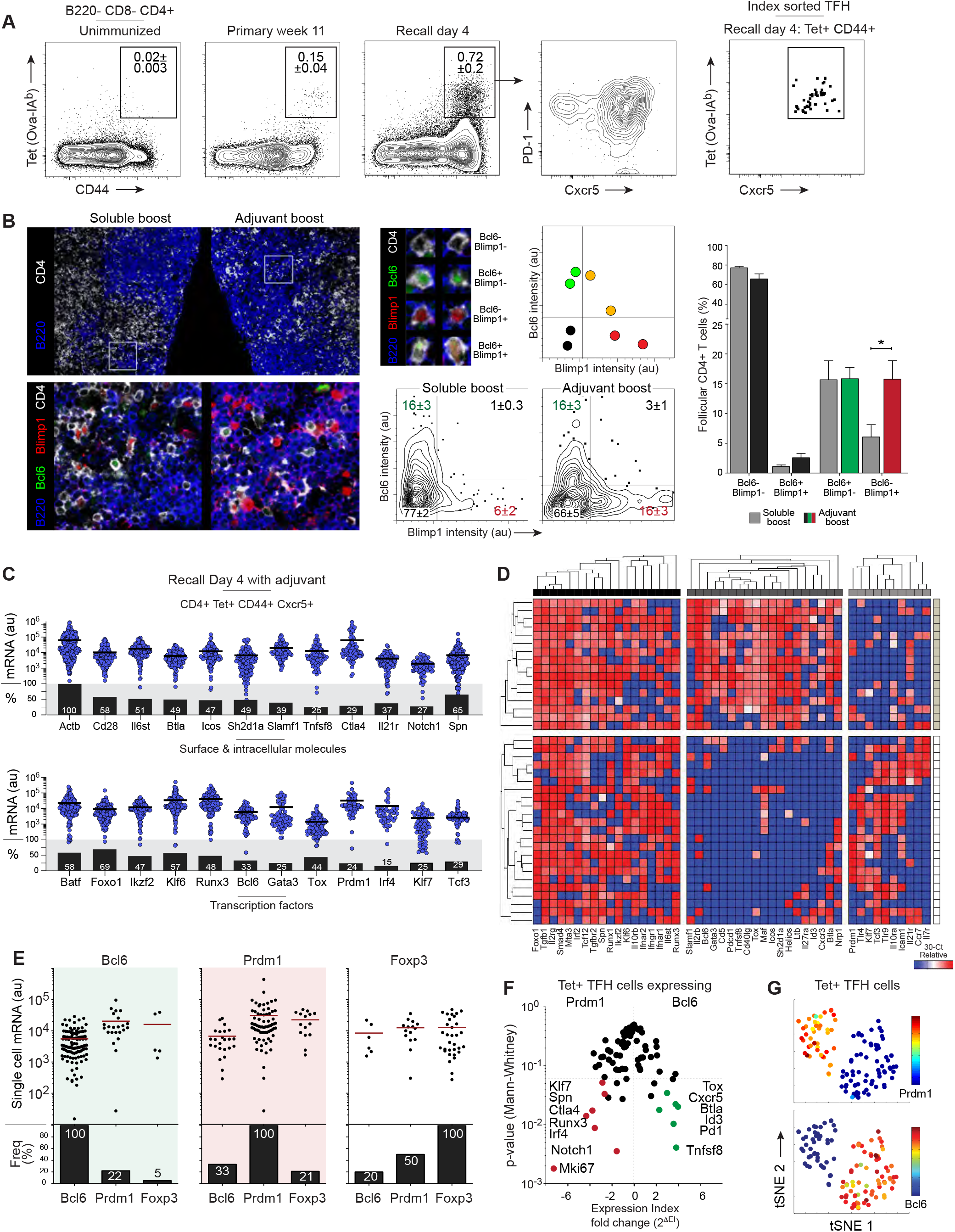
Antigen-specific memory PC T_FH_ and GC T_FH_ responders. C57BL/6 mice were immunized subcutaneously at the base of the tail with 400μg NP-Ova in MPL adjuvant. At >10 weeks post-primary immunization, secondary immunization was performed with 100μg NP-Ova with and without adjuvant MPL. Mice were sacrificed at day 4 post-secondary immunization. **A)** Ova-IA^b^ tetramer labeling of CD4^+^ draining LN cells in unimmunized (n=3 mice), 11 weeks post-primary (n=3 mice) or 4 days post-antigen recall with adjuvant (left) (n=6 mice from 4 experiments), a flow plot display of PD1 and CXCR5 expression within Ova-IA^b+^ T cells (center) and index-sorted Cxcr5^+^Ova-IA^b+^ T_FH_ sorted for multiplexed single-cell qPCR analysis (right). **B)** Confocal microscopy of individual lymph nodes post-antigen recall with adjuvant or soluble boost displaying CD4 (white), B220 (blue), Bcl6 (green) and Blimp1 (red) expression (left), quantification of the confocal image intensity of Bcl6^-^Blimp1^-^ (black), Bcl6^+^Blimp1^-^ (green), Bcl6^+^Blimp1^+^ (orange) and Bcl6^-^Blimp1^+^ (red) subsets (center) and bar graph display of Bcl6 and Blimp1 subsets with soluble (grey) or adjuvant (black, green, or red) boost. **C)** Single-cell multiplexed qPCR was performed on antigen-specific day 4 post-recall T_FH_ (B220^-^CD8^-^CD4^+^CD44^+^Ova-IA^b+^CXCR5^+^) sorted in (A). Dot plot display of mRNA expression of surface and intracellular molecules (top) or transcription factors (bottom) from single OVA-IA^b^ tetramer binding memory T_FH_ cells and bar graph displays of population penetrance within the total recall T_FH_ compartment (n=296). Each blue dot represents a single cell, with the geometric mean indicated as a black line. **D)** Hierarchical clustering of single qPCR gene expression according to gene expression values (Ct) by average linkage hierarchical clustering (Euclidian distance) and displayed red (highest) to blue (lowest). Each row depicts expression within a single cell (n=39) and each column depicts a single gene. Clusters were generated using GenePattern. K-means clusters are depicted for cells and genes by heatmap (n=2 mice). **E)** A dot plot display of single Ova-IA^b+^ T_FH_ cells expressing *Bcl6* (green, n=115), *Prdm1* (red, n=73) or *Foxp3* (white, n=33) depicting mRNA expression (top) and bar graph display of frequency of each gene (bottom). Each black dot represents a single cell, with horizontal bars indicating geometric mean. Black bar graphs display frequency of each selected gene. **F)** A volcano plot display of differential gene expression between *Prdm1*^+^T_FH_ (upregulated in red) and *Bcl6*^+^T_FH_ cells (upregulated in green) and **G)** tSNE dimensionality reduction of gene expression (30-Ct) from *Prdm1*^+^ or *Bcl6*^+^ T_FH_ cells. tSNE map overlay depicts *Prdm1* expression (top) or *Bcl6* expression (bottom) for individual T_FH_ cells, color coded from dark blue (no expression) to dark red (high). Single cell qPCR analysis (C-G) included post-QC analysis of selected genes (n=82) on antigen-specific CXCR5^+^ T-cells (n=296) from four independent experiments with 1-2 mice per experiment.

For gene expression studies, we chose the Biomark platform that targets 96 genes for amplification and uses qPCR for quantification. As with previous qtSEQ analysis, we used cell indexing to directly sort rare populations of antigen-specific T_FH_ compartments and included a nested PCR intron targeting for each assay in library formation to increase the sensitivity for each assay. The high sensitivity and penetrance frequencies are displayed as calculated arbitrary units of mRNA for a representative set of target genes with estimates for the proportion of positives expressing each signal (**Fig. 6C**). Unsupervised K means clusters predicting three divisions by two subsets broadly segregates a set of high penetrance shared gene expression with separation of mRNA species associated with *Bcl6* expressing GC-T_FH_ and a set associated with *Prdm1* expressing PC-T_FH_ (**Fig. 6D**).

Due to recent antigen-specific activation there is high expression and frequency for most of the shared genes across both sub-populations as shown in the example. A series of commonly expressed cytokine receptors and surface molecules (eg *Il2rg, Tgfbr2, Il10rb, Ifnar2, Ifnar1, Il6st*, *Spn*) accompany multiple transcription factors known to influence lymphocyte differentiation (eg *Foxo1, Smad4, Mta3, Irf2, Tcf12, Runx1, Runx3, Ikzf2 and Klf6*). The *Bcl6* associated GC-T_FH_ cluster differentially expresses many surface (eg *Slamf1, Ilr2b, Cd5, Pdcd1, Tnfsf8, Cd40l, Icos, Cxcr3, Btla, Nrp1*) and intracellular/secreted molecules (eg *Sh2d1a, Ltb*) capable of modifying B cell differentiation. Many of the co-expressed transcriptional regulators (eg *Gata3, Tox, Maf, Helios, Id3*) are well known to influence T_H_ cell function and GC B cell differentiation. While less represented, multiple gene products assorted with the *Prdm1* associated cluster (eg *Tlr4, Il10ra, Icam1, Il21r, Ccr7, Il7r*) with transcription factors *Klf7* and *Tcf3* being the most closely co-expressed regulators provide a glimpse of the differential program of an antigen-activated memory-response PC-T_FH_ compartment.

Across multiple experiments, we accumulated many examples of pMHCII^+^ T_FH_ cells that expressed the major regulators *Bcl6, Prdm1* and *Foxp3* (**Fig. 6E**). The vast majority of *Bcl6* expressing GC-T_FH_ cells at this timepoint did not co-express *Prdm1* or *Foxp3*. Similarly, the majority of *Prdm1* expressing PC-T_FH_ did not express *Bcl6 or Foxp3*. However, half of the *Foxp3* expressing T_FR_ cells co-expressed *Prdm1* in a manner resembling the Blimp1^+^ T_FR_ regulatory compartment seen at steady-state. Unlike typical T_FR_ compartments known to be self-reactive, these Blimp1^+^ T_FR_ cells are sorted based on shared antigen-specificity. This latter compartment could be considered putative PC-T_FR_ cells that negatively impact memory-response reactivation to select higher affinity memory B cells into PC differentiation at recall.

Focusing analysis on the GC-T_FH_ to PC-T_FH_ transcriptional division, we compared the gene expression across these compartments directly. The volcano plot displays a set of significant differences with antigen-specific GC-T_FH_ expressing higher levels of multiple genes associated with the GC pathway (eg *Cxcr5, Pdcd1, Tox, Id3, Btla, Tnfsf8*) contrasting with antigen-specific *Prdm1*^+^ PC-T_FH_ differential surface expression of *Spn* (CD43), *Ctl4a* and transcriptional regulators *Klf7, Irf4*, and *Runx3* (**Fig. 6F**). Using these key differences, dimensionality reduction separates these compartments into separate regions to segregate these single antigen-reactive cells across the GC-T_FH_ and PC-T_FH_ compartments (**Fig. 6G**). These results demonstrate the rapid emergence of divergent follicular pathways with evidence for antigen-specific Blimp1 expressing T_FR_ cells and separate pathways of Bcl6^+^ GC-T_FH_ and Blimp1^+^ PC-T_FH_ at antigen recall.

## DISCUSSION

These studies demonstrate an early cellular bifurcation in follicular T cell differentiation that regulates the dichotomous antigen-driven PC and GC B cell fate. Separable persistent PC-T_FH_ and GC-T_FH_ compartments exist at homeostasis based primarily on the reciprocal expression of major transcriptional repressors Blimp1 and Bcl6. Conditional deletion of *Prdm1* in mature CD4 T cells significantly compromised early PC differentiation with no impact on GC formation. Furthermore, *Prdm1* expression was needed to support significant post-GC PC differentiation indicating the need for persistent, separable and potentially GC-localized PC-T_FH_ function. Thus, Blimp1 is a key transcriptional mediator of PC-T_FH_ function and inter-dependence of both T_FH_ subset functions is crucial for the generation of high-affinity long-lasting B cell immunity.

The central role of Bcl6 within both B cells (*1–3*) and GC-T_FH_ cells (*4–6*) is well established and required to promote and support antigen-specific GC B cell function. The central GC function of BCR affinity maturation creates long-lived memory B cell and post-GC PC compartments needed for durable immune protection (*1, 2*). GC-T_FH_ orchestrate this cycle of clonal selection (*2, 38*) impacting B cell outcomes through dynamic and varied inter-cellular cognate contact (*6, 38, 45-47*). The molecular rules of cellular exit from the GC reaction are less well understood (*2, 48, 49*). Quantitative differences in T_FH_-GC B cell contact duration and quality play a role (*6, 50*). B cell transcriptional changes occur to down-regulate Bcl6 and permit alternate programs and outcomes. One dominant outcome is the upregulation of Blimp1 and the elaborate re-programming of post-GC PC differentiation (*40, 51*). Here, we propose that cognate contact with PC-T_FH_, either within the GC or upon GC exit, is required to imprint this terminal differentiation program and evoke continued secretion of high-affinity antibody.

Multi-targeted transcriptional repression exerts its function in cell-type and differentiation-stage specific ways (*4, 12*). As described, CD4-expressed *Prdm1* is highly heterogenous in its impact on effector and regulatory T cell functions with influence during early T cell development (*25, 26, 40*). Here, we identify multiple sub-compartments of Blimp1^+^ PC-T_FH_ and elements of a differentially-expressed transcriptional program. The steady-state splenic PC-T_FH_ program diverged significantly from the GC-T_FH_ compartment deviating away from multiple facets of a GC-B cell inducing program (*12, 52, 53*). It was important for these initial studies to segregate cellular subsets based on persistent expression of the central factors Blimp1 and Foxp3. Multiple surface and nuclear expressed molecules segregated into the PC-T_FH_ compartment to establish elements of PC-enhancing functional potential in the steady-state. It remains important to resolve which components of the PC-T_FH_ molecular machinery support the spectrum of central PC functions necessary for rapid antibody production, PC terminal differentiation, long-term niche survival, continued secretion of antibodies and ultimately induce antigen-specific memory PC formation at recall.

Regulatory T cell compartments counterbalance helper T cell activities in both conventional and follicular effector T cell functions. Foxp3 expressing T_FR_ cells impose a negative influence on the T_FH_ spectrum of GC B cell inducing functions (*31, 32, 54*) and indirectly on post-GC PC differentiation (*29, 30*). Here, we demonstrate that Blimp1 expression also segregates the CXCR5^+^PD1^hi^ T_FR_ compartment and speculate that the distribution of this known T_FR_ associated molecule differentially associates with PC-regulating functions. Surprisingly, we also detected large fractions of GITR^+^Blimp1^+^ T_FH_ that did not express Foxp3 in the steady-state. Gene expression analysis of these two compartments indicates the presence of multiple immune checkpoint inhibitors with less effector enhancing capacity, consistent with a negative regulatory function for both subsets (*31*). Furthermore, antigen specificity of T_FR_ action remains controversial (*34*) with recent evidence for the conversion of GC-T_FH_ to GC-T_FR_ (*35*). Here, foreign antigen-specificity is shared across memory T_FR_ subsets of both Foxp3^+^Bcl6^+^ GC-T_FR_ and Foxp3^+^Bcl6^-^ Blimp1^+^ T_FR_ that may be limiting PC differentiation as an early outcome of the GC reaction. Based on the distribution of Blimp1 and Bcl6, we propose the existence of separable Foxp3^+^ GC-T_FR_ and PC-T_FR_ follicular regulatory compartments.

These follicular helper sub-compartments are central to multiple aspects of adaptive immunity with relevance to recent reports of adaptive immune tolerance, the tumor microenvironment, infectious disease and vaccine responses. We recently reported the impact of PD1 blockade on adaptive immune tolerance that selectively dysregulates a type 2 GC-T_FH_ program (*42*) and exaggerates IgG1 GC B cell expansion and excessive amounts commensal reactive antibodies that may involve separate levels of regulation (*43*). Recent reports indicate that T_FH_ cells and B cells in the tumor microenvironment are also key to immune checkpoint therapy indicating another broad application of our current findings in this clinical setting (*55*). The range and dynamics of the immune response in humans with COVID-19 has been probed at great depth with a spectrum of CD4 T_H_ attributes driving B cell helper functions that correlate with protection (*56–59*). Some trends indicate regulatory CD4 involvement (*60*) and correlations of circulating T_FH_ cells with convalescence or expression of mild disease in this infection (*61, 62*). In acute COVID-19, extrafollicular B cell responses alone were insufficient for protection (*20*) and the T_FH_ compartment can be compromised upon infection leading to decreased GC B cell formation and lower quality antibody responses (*21*). SARS-CoV2 mRNA vaccines induce increased presence of local T_FH_ and GC B cell responses that correlate with immunity (*63*). Similar correlations are found in malaria infection (*64*), influenza vaccination (*65*) and long-lived T_FH_ compartments that support humoral immunity (*66*). Thus, a division of function in these central and potentially inter-dependent foreign antigen-driven T_FH_ functions must be considered across multiple facets of follicular T cell controlled B cell adaptive tolerance, tumor development and immunity to infection.

While PC-T_FH_ cells alone are required for early and rapid antibody responses, both T_FH_ sub-classes are essential for generation of high-affinity long-lived PC compartments. Recent studies demonstrate non-T_FH_ cells lacking Bcl6 that drive class-switch and PC differentiation (*19*). It is plausible that Blimp1 expressing ETH localizing close to the T-B borders can direct PC fate or alternatively these Bcl6^-^ T_H_ cells more closely resemble Blimp1^+^ PC-T_FH_ described here. Specialized GC-T_FH_ subsets are also reported with functions that selectively regulate isotype-specific B cell responses (*37, 42, 67, 68*). It is of interest to speculate whether these divided traits extend into the PC-T_FH_ compartment. Selectively targeting memory PC-T_FH_ compartments also provides the means to bias memory B cell fate at antigen recall to enhance long-term immunity. Cellular organization and molecular components of PC-T_FH_ transcriptional program indicate functional sub-specialization with pathways that can be separately targeted for immunotherapeutic purposes and adjuvant design in future vaccines.

## ACKNOWLEGMENTS

BioRender.com was used to construct the graphical summary.

## Funding

This work was supported by the US National Institutes of Health (AI047231, AI040215 and AI071182) and Bill & Melinda Gates Foundation (BMGF 0PP1154835) to M.G.M.-W. P.J.M received a Young Investigator Award from the Bettencourt-Schueller Foundation.

## Author contributions

K.B.M, L.M.W and M.M.W conceived the study. K.B.M, A.G.S, B.W.H, S.M.S, P.J.M, L.M.W and M.M.W designed and performed experiments and analyzed, interpreted data. K.B.M, L.M.W and M.M.W wrote the manuscript.

## Competing interests

There are no competing interests.

## Materials and Methods

### Mice

C57BL/6, B6.SJL-Ptprc^a^Pepc^b^/Boy (congenic C57BL/6-Ly5.1, CD45.1^+^), B10.BR, B6.129S7-Rag1^tm1Mom^ (Rag1^def^) (Jackson Laboratory) were bred and housed at The Scripps Research Institute. Prdm1^f/f^Cre-ER^T2^ were bred by backcrossing C57BL/6-*Prdm1*^flox/flox^ (*51*) (from Kathryn Calame, Columbia University) and C57BL/6-Et(cre/ERT2) for >12 generations. Blimp1-YFP (*Prdm1*-YFP) mice (*27*) (provided by Susan Kaech, Salk Institute) and Foxp3-IRES^Thy1.1^ reporter mice (*69*) (provided by Ye Zheng, Salk Institute) were bred to generate dual Blimp1-YFP/Foxp3-Thy1.1 reporter mice. Male and female mice aged 8-20 weeks were used in this study. All mice were bred and maintained under specific pathogen-free conditions. Procedures were performed in accordance with protocols approved by The Scripps Research Institute’s Institutional Animal Care and Use Committee (IACUC).

### Immunizations

The hapten 4-hydroxy-3-nitrophenylacetyl (NP) (Biosearch) was conjugated to carrier proteins keyhole limpet hemocyanin (KLH) (Thermo Fischer), pigeon cytochrome C (PCC) (Pierce) or ovalbumin (OVA) (Thermo Fischer). Mice were immunized subcutaneously at the base of the tail or intraperitoneal injection with 400μg of hapten-carrier conjugate mixed with monophosphoryl lipid A (MPL)-based adjuvant, as indicated. This oil-in-water adjuvant was formulated in-house by generating an emulsification with MPL (Avanti), lecithin (Spectrum) and Tween-80 (Sigma) in squalene oil (Sigma) (*44*).

### Flow cytometry and cell sorting

Single cell suspensions of spleen, lymph node, or femur bone marrow were prepared and processed as previously described (*44*). For intracellular staining, cells were additionally incubated with eFluor780 viability dye (ThermoFisher) for 15 minutes and then processed using the Foxp3 staining kit (ThermoFisher) per manufacturer’s protocol. Cells were then stained for transcription factors Blimp1 (BD Biosciences) or Foxp3 (eBioscience) for 30 minutes. All flow cytometry was performed on a four-laser FACS Aria III using Diva 8.0 software (BD). Downstream analyses, including index sort plate-mapping and tSNE visualization, were performed using FlowJo software (Version 10, TreeStar).

### Adoptive transfer and tamoxifen administration

Single-cell suspensions of 2.5x10^6^ CD4^+^ T cells and 1x10^7^ CD19^+^ B cells were sorted separately via flow cytometry from Prdm1^f/f^Cre-ER^T2^ spleens. CD4^+^ T cells were then transferred to complete RPMI media (10% fetal bovine serum) and treated with 1μM 4-hydroxytamoxifen (Sigma) freshly diluted in 95% ethanol for 1 hour at 37°C. CD4^+^ T cells were washed 3 times in complete RPMI media, combined with the untreated CD19^+^ B cells and transferred through intraperitoneal injection into Rag1^def-^ recipient mice.

### In vitro co-cultures

Splenic CD4^+^ T cells were isolated from Blimp1-YFP reporter mice and separated using flow cytometry based on phenotype. Twenty thousand activated T cells were combined with sixty thousand naïve T feeder cells in 96 well plates with 1μg/mL α-CD3ε (Bio X Cell), 2.5μg/mL CD28 (Bio X Cell) and 10U/mL IL-2 (Sigma) in complete RPMI. After 48 hours, T cells were replated into new wells. At 96 hours, 3x10^5^ CD19^+^ B cells were added to pre-existing wells containing T cells and supplemented with 200ng/mL BAFF (Sigma) in complete RPMI for 5 days. Plasma cells were analyzed by flow cytometry and Elispot.

### ELISPOT assay

Multiscreen HTS plates were coated with IgM, total IgG or IgG1 (BioLegend) at 4°C overnight. Cells were added to each well and incubated at 37°C. After 18 hours, wells were incubated with the secondary antibody conjugated to horse radish peroxidase (Southern Biotech) and developed using a filtered solution of 0.2mg/mL 3-amino-9-ethylcarbazole (Sigma) and 0.015% H_2_O_2_ (Sigma) in 0.1M sodium acetate (pH 4.8-5.0, Sigma) for 3-8 minutes. Wells were washed with H_2_O and allowed to dry at least 48 hours before visual inspection of spots and manual enumeration using a stereomicroscope.

### Single cell real-time PCR assay

Lymph nodes were collected and stained for flow cytometry. For tetramer staining, cells were stained for 45 minutes at room temperature with PE-OVA-IA^b^ (*70*) (from Luc Teyton, Scripps Research). Single T cells were then sorted into 96-well plates and processed as previously described (*44*). To confirm fidelity of sorting and processing, at least four wells per plate had no cells sorted, but were fully processed with the plate as a negative control. Following reverse transcription, cells were processed using the BioMark Real-time PCR system (Fluidigm), TaqMan Universal PCR Master Mix (Applied Biosystems) and a curated TaqMan gene expression assay probing ∼90 genes (*44*). Cells that did not express at least two reference genes (Actb, Gapdh, B2m) were removed from subsequent analyses (<5% of cells). Cycling threshold (Ct) values from these individual cells were transformed into relative mRNA abundance through subtraction of the Ct value from the baseline of 30 and subsequent conversion to a numerical value at 2(30−Ct) for display on a log scale.

### Microscopy

Draining LNs were fixed overnight at 4 °C in paraformaldehyde-lysine-periodate buffer, dehydrated in 30% sucrose, embedded in OCT medium (Tissue-Tek) and frozen on dry ice. Tissues sections were sliced into 8μm sections and processed as previously described (*44*). Tissues were stained with α-B220 (BioLegend), α-CD4 (GK1.5), α-Blimp1 (Santa Cruz) and α-Bcl6 (BD Biosciences). Slides were examined using the Nikon C2 confocal microscope with four lasers (408 nm, 488 nm, 543 nm and 633 nm) and a 60× objective. Images were acquired with Nikon Elements software and were further processed with Adobe Photoshop.

### Single-cell index sort, library preparation, and RNA Sequencing (qtSEQ)

T cells were sorted from single Blimp1-YFP reporter or dual Blimp-YFP/Foxp3-Thy1.1 spleens into 384-well plates, with index sort information collected on each cell: naive (CD8^-^B220^-^ CD4^+^TCRβ^+^ CD62L^+^CD44^-^ Thy1.1^-^), GC TFH (CD8^-^B220^-^ CD4^+^TCRβ^+^ CXCR5^+^GITR^-^Blimp1^-^Thy1.1^-^), PC TFH (CD8^-^B220^-^ CD4^+^TCRβ^+^ CXCR5^+^GITR^-^Blimp1^+^Thy1.1^-^), GITR^+^ PC TFH (CD8^-^B220^-^ CD4^+^TCRβ^+^ CXCR5^+^GITR^+^ Blimp1^+^Thy1.1^-^) and Blimp1+ TFH (CD8^-^B220^-^ CD4^+^TCRβ^+^ CXCR5^+^GITR^+^ Blimp1^+^Thy1.1^+^) for dual reporter sorts. Naïve CD4 T cells (CD8^-^B220^-^ CD4^+^TCRβ^+^ CD62L^+^CD44^-^), GC TFH (CD8^-^B220^-^ CD4^+^TCRβ^+^ CXCR5^+^GITR^-^Blimp1^-^), PC TFH (CD8^-^B220^-^ CD4^+^TCRβ^+^ CXCR5^+^GITR^-^Blimp1^+^), ETH (CD8^-^B220^-^ CD4^+^TCRβ^+^ CXCR5^-^GITR^-^Blimp1^-^) and Blimp1^+^ ETH (CD8^-^B220^-^ CD4^+^ TCRβ^+^ CXCR5^-^GITR^-^Blimp1^+^) for single Blimp1YFP reporter sorts. Doublets were excluded using SSC-A and SSC-H parameters.

Transcriptional analysis of single cells was carried out using a custom RNA sequencing protocol, quantitative and targeted single cell sequencing (qtSEQ) (*39*). As previously described, this technique sequences polyA mRNA using a nested set of 3’ targeted primers to amplify specific gene (*37, 42, 43, 71*). Final cDNA amplified library concentration and size was determined using the Qubit 4 Fluorometer (ThermoFisher) and 2100 BioAnalyzer (Agilent). Multiplexed libraries were sequenced with 1% PhiX control library on the Illumina NextSeq 500 per manufacturer’s protocol (Illumina, Read 1: 19 cycles; Index Read 1: 6 cycles; Read 2: 67 cycles).

### qtSEQ data processing and quality control

Illumina base call (bcl) files were converted to FASTQ files using bcl2fastq software (Illumina). FASTQ files were then demultiplexed using a barcode demultiplexing script adapted from Cel-Seq2 (*72*). Well barcodes with a quality score **o**f <Q10 were removed. Data was aligned with Bowtie2 (v2.2.9) to a custom genome, consisting of amplicon regions of specifically targeted genes based on murine genome GRCm38.p6, version R97 (Ensembl). Read and UMI tabulation reduction were performed using HTSeq (*73*) and any duplicate UMIs were removed from all downstream analyses. Index sort information analyzed and exported from FlowJo (v10, Treestar) was manually aligned with each corresponding well.

Analysis of single-reporter Blimp1-YFP T splenocytes included 1425 total cells across two experiments from 2 mice and dual reporter Blimp1-YFP/Foxp3-Thy1.1 studies included 1204 total cells across two experiments from 7 mice. Each T cell contained at least one count of *Cd3d*, *Cd3e* or *Cd3g* and cells with less than 60 UMIs, more than 2000 UMIs or less than 25 unique genes detected were excluded from downstream analyses. Data was pooled into a single count matrix and processed using the R package Seurat (v3.2.3) (*74*). Scaling and normalization were performed using the R package sctransform (v0.3.2.9002) (*75*), incorporating the batch_var function to account for variation between experiments.

### scRNA-seq computational analysis

Differential gene expression was calculated using the R-package MAST (Model-based Analysis of Single Cell Transcriptiomics, v1.16.0) (*76*) to calculate fold change and p-values for all genes. Heatmaps of pseudobulk populations were generated within Seurat using the “DoHeatmap” function. The R package pheatmap (v.1.0.12) was utilized to generate heatmaps including euclidean distance hierarchical clustering based on averaged UMI count per gene. Dot plots were generated in Seurat using the “DotPlot” function. Uniform Manifold Approximation and Projection (UMAP) was performed in Seurat, clustering with manhattan metrics to determine nearest neighbors. All volcano plots were created using Prism software (GraphPad, v9.0).

Network analysis was generated using the web application Metascape (*41*). Lists of differentially or uniquely upregulated genes from GC-T_FH_ or PC-T_FH_ relative to naïve T cells were input and densely connected networks were highlighted by the molecular complex detection algorithm (MCODE).

### Statistical analysis

Mean values, s.e.m. values, unpaired t-tests and Mann-Whitney tests were calculated and graphed with Prism software (GraphPad). A P value of less than 0.05 was considered statistically significant.

---

**Fig. S1.**
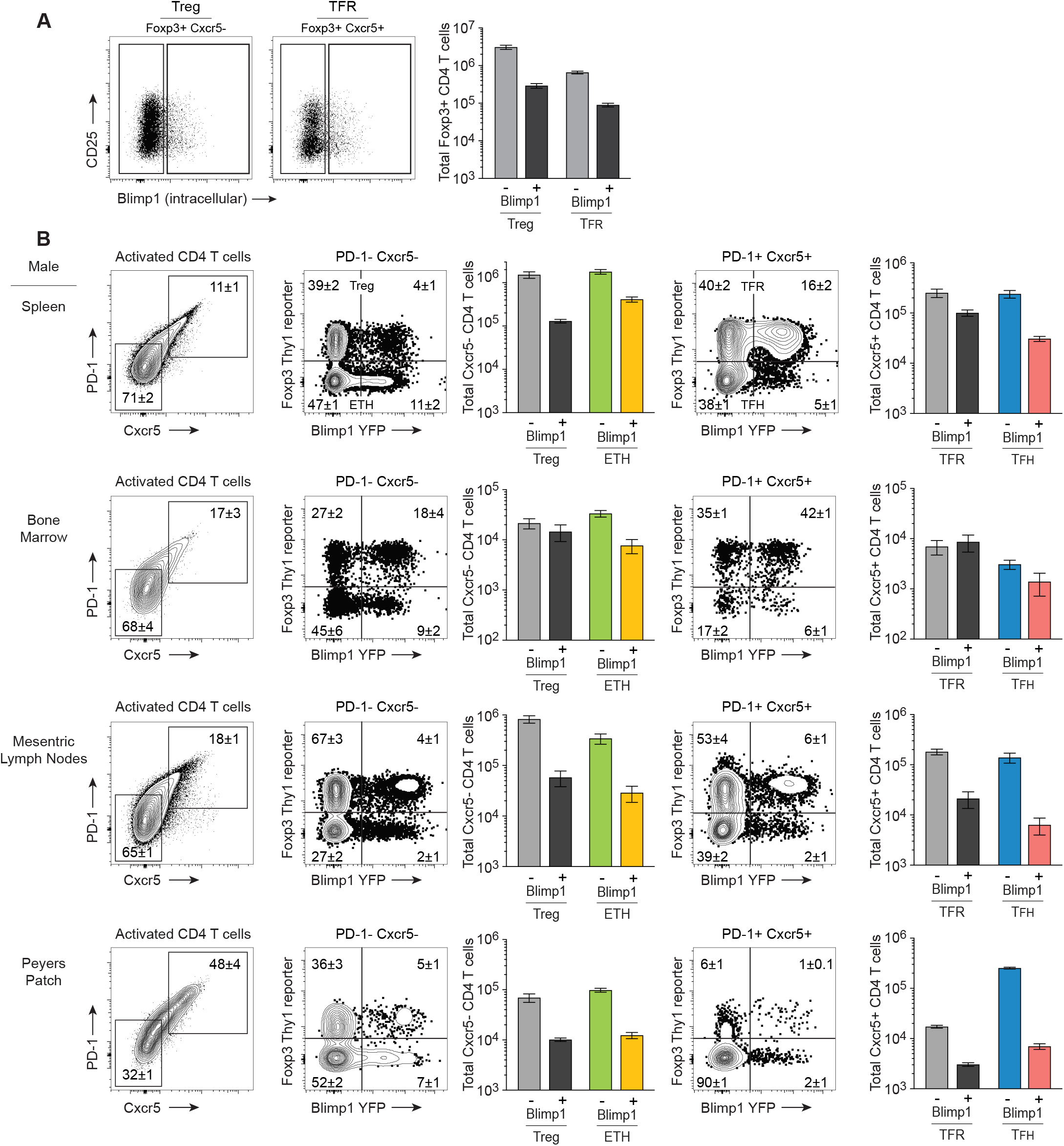
Blimp-1 expressing TFH subsets exist across multiple lymphoid tissues. **A)** Flow cytometry analysis of CD25 and Blimp1 distribution within dual Blimp1-YFP/Foxp3-Thy1.1 reporter regulatory T splenocytes (B220-CD8-TCRb+CD4+ CD44+GITR+Foxp3+). **B)** Multi-organ analysis of Blimp1 and Foxp3 distribution within PD-1-CXCR5-non-follicular or PD-1+ CXCR5+ follicular compartments of male dual Blimp1-YFP/Foxp3-Thy1.1 reporter mice using flow cytometry. Numbers in flow cytometry plots display mean±SEM (n=4).

**Fig. S2.**
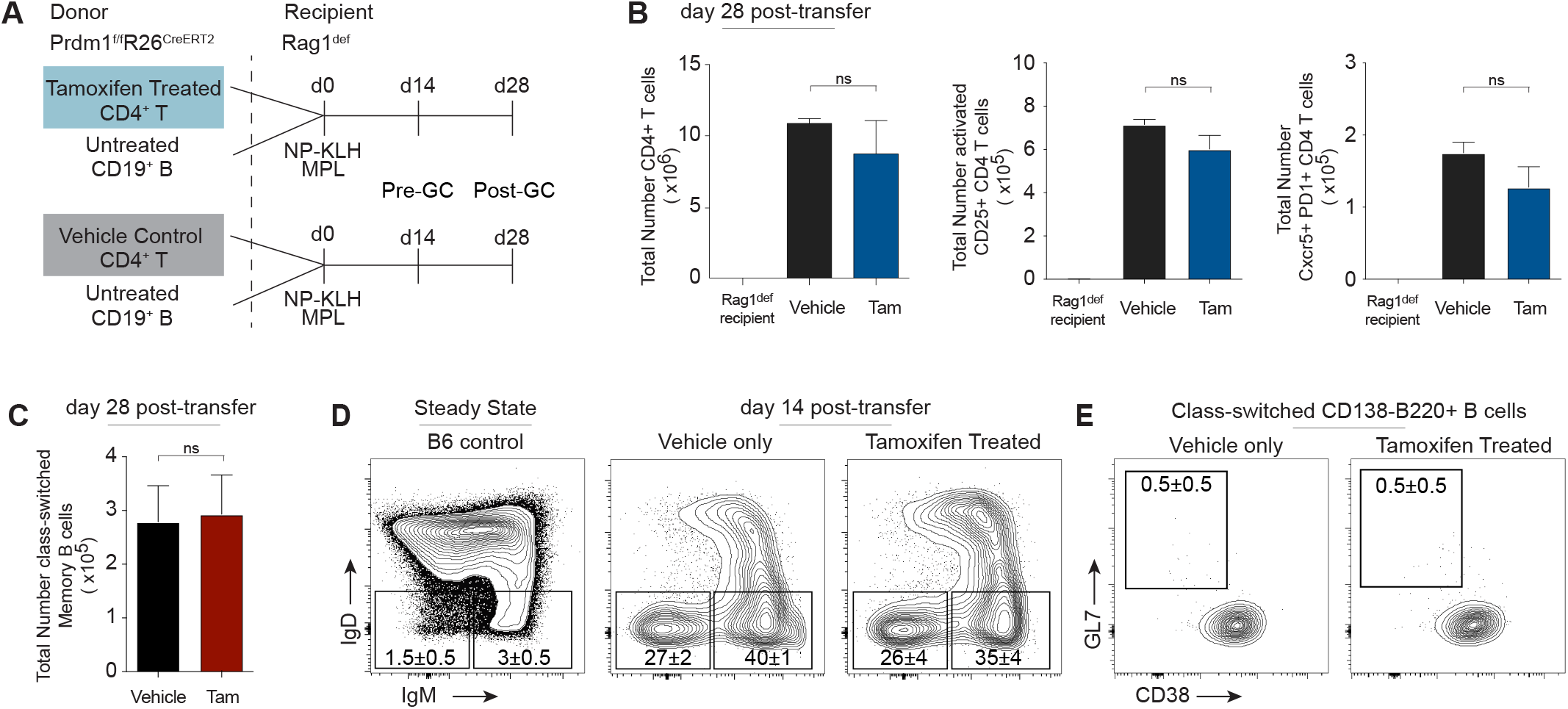
Blimp-1 deleted CD4 T cells in adoptive transfers. Adoptive transfer of Prdm1^fl/fl^ CD4+ T splenocytes treated with or without tamoxifen (2.5x10^6^) and untreated B cells (1x10^7^) transferred i.p. into Rag1^def^ recipients and immunized with 400μg NP-KLH in adjuvant. Mice were sacrificed and analyzed on day 14 or 28. **A)** Experimental design of adoptive transfer experiments. **B)** Total cell numbers of CD4+ T cells, activated CD25+ T cells, activated Cxcr5+PD-1+ or **C)** B220+GL7-CD38+ memory B cells on day 28 post-transfer, depicted as bar graphs [grey, Rag1^def^ recipient control; black, vehicle (T, B cells); blue, tamoxifen-treated (T cells); red, tamoxifen-treated (B cells)]. **D)** Representative flow plot display of total B cells (Gr1-CD3-CD138+ and/or CD19+) in steady state C57BL/6 control (left), vehicle (center) and tamoxifen-treated (right), gating on IgM (IgD-IgM+) and class-switched (IgD-IgM-) subsets and **E)** GC B cells (GL7+CD38-) from the class-switched subset (IgD-IgM-B220+CD138-) on day 14 post-adoptive transfer. Student’s t test: ns= not significant. Flow plots display mean±SEM. Data are representative of three independent experiments with n= 6-7 total mice per timepoint.

**Fig. S3.**
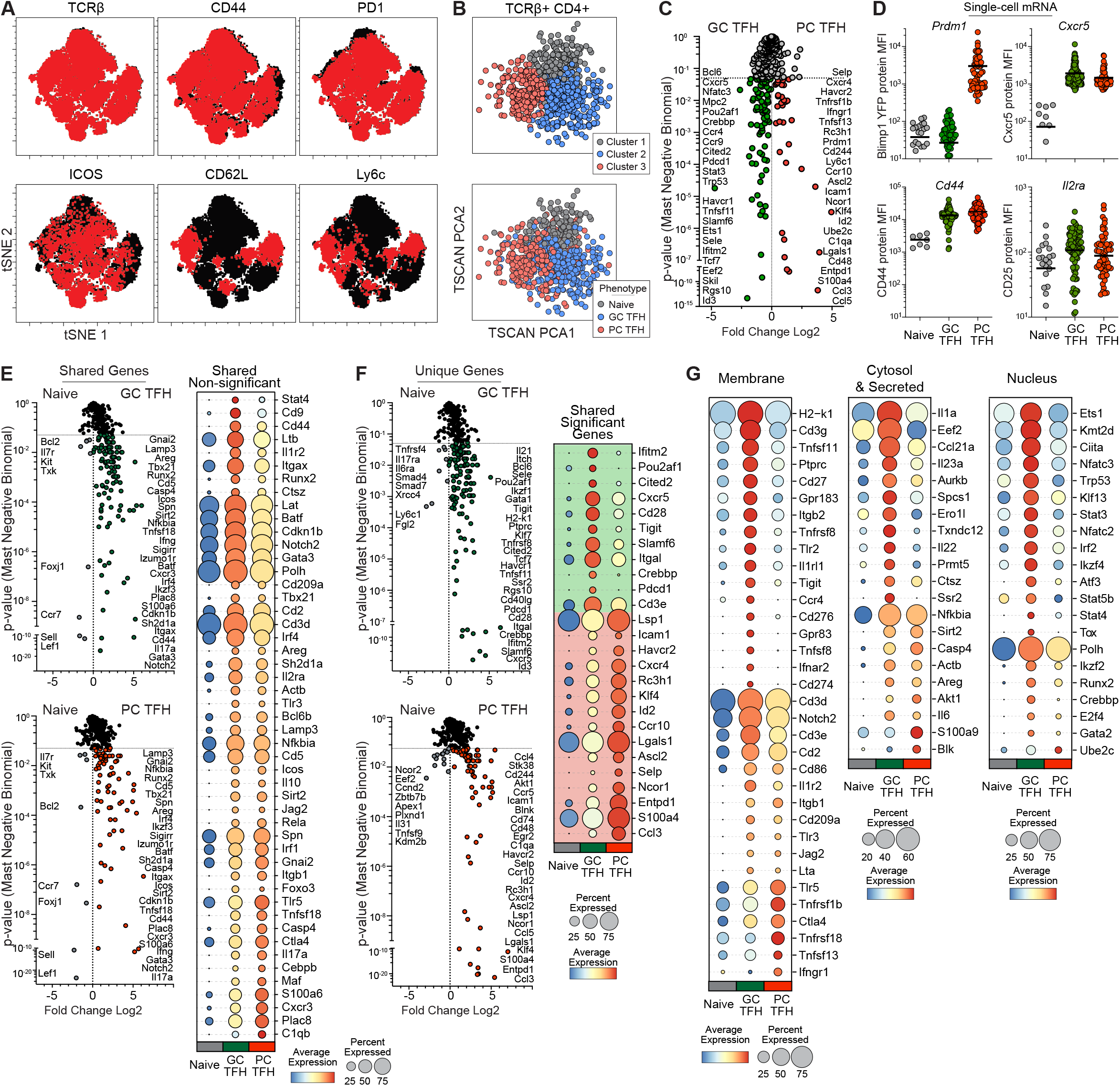
Shared and unique components of the PC TFH and GC TFH programs. **A)** tSNE dimensionality reduction of Cxcr5+ TFH splenocytes from unimmunized dual Blimp1-YFP/Foxp3-Thy1.1 reporter mice (black) overlayed with protein expression from individual cells (red). **B)** Single cell RNAseq (qtSEQ) on naive T, GC TFH and PC TFH index sorted populations. TSCAN pseudotime alignment of three sorted subsets as independently clustered (top) and overlayed with index-sorted phenotype (bottom). **C)** Volcano plot comparing differential gene expression between GC TFH (upregulated in green) and PC TFH (upregulated in red). **D)** Analysis of protein Mean Fluorescent Intensity (MFI) from index sorted cells with corresponding single cell RNA expression. **E)** Differential gene expression between naïve and GC TFH or naïve and PC TFH displaying significantly upregulated genes that are shared between both GC TFH and PC TFH or **F)** uniquely upregulated in only GC TFH or PC TFH or significantly different, but shared upregulation, as depicted by volcano plots (left) or dot plots (right). **G)** A dot plot display of significantly differentially expressed genes within GC TFH, PC TFH, or naïve, organized by population and frequency. Dot color reflects average gene expression and dot size depicts percent expressed. Single cell RNAseq analysis (qtSEQ; B-G) included post-QC subsets of GC TFH (n=304), PC TFH (n=248) and naïve T (n=153) from two independent experiments with 3-4 mice per experiment.

**Fig. S4.**
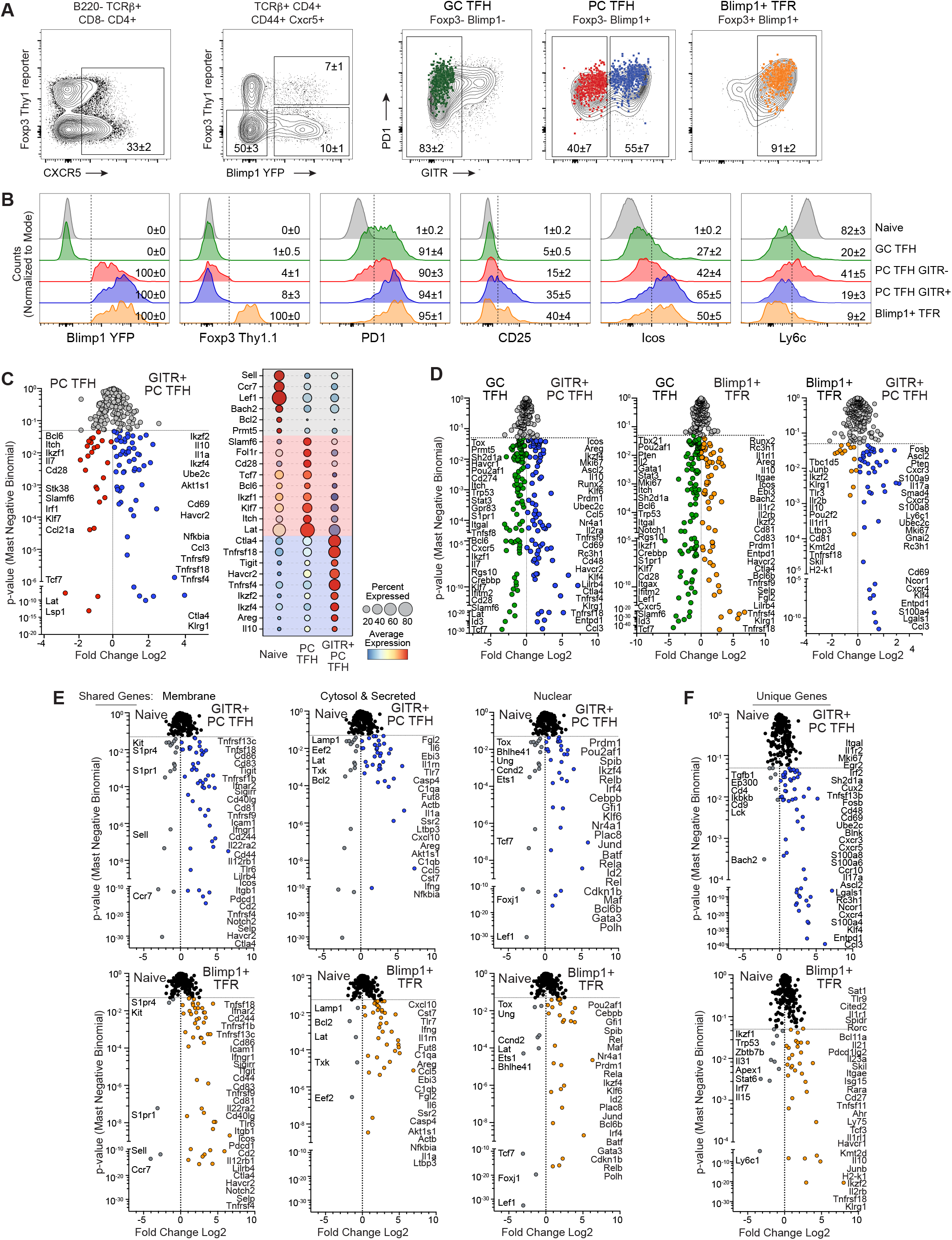
CXCR5+Blimp1 subsets contain transcriptionally distinct enhancers and regulators. Steady state splenocytes were sorted for single cell RNAseq analysis from activated CXCR5+ subsets. **A)** Index sort of follicular T (B220-CD8-TCRβ+CD4+CD44+Cxcr5+) subsets displaying PD-1 and GITR expression within GC TFH (Blimp1-GITR-Thy1.1-), PC TFH (Blimp1+GITR+/-Thy1.1-) and Blimp1+ TFR (Blimp1+GITR+Thy1.1+) and **B)** differential protein expression within sorted subsets displayed as histograms. **C)** Differential gene expression from single-cell RNAseq (qtSEQ) analysis on PC TFH comparing GITR+ and GITR-subsets displayed in (A) as a volcano plot (left) and dot plot (right). Dot color reflects average gene expression and dot size depicts population frequency. **D)** Volcano display of differential gene expression within sorted TFH subsets GC TFH and GITR+ PC TFH (left), GC TFH and Blimp1+ TFR (center) and Blimp1+ TFR and GITR+ PC TFH (right) sorted in (A). **E)** A volcano display of differential gene expression in Blimp1+ TFR to naive and GITR+ PC TFH to naïve depicting shared genes upregulated in both data sets and **F)** genes uniquely upregulated in GITR+ PC TFH (top) or Blimp1+ TFR (bottom). Numbers in flow plots display mean±SEM (n=8). Single cell RNAseq analysis (qtSEQ); B-E) included post-QC subsets of GC TFH (n=304), PC TFH (n=248) and naïve T (n=153) from two independent experiments with 3-4 mice per experiment.

## Notes

### Competing Interest Statement

The authors have declared no competing interest.

